# Varied mutual growth inhibition between commensal yeasts and *E. coli* strains

**DOI:** 10.1101/2025.05.29.656793

**Authors:** Eugénie Pingault--Ordronneau, Antoine François, Vashineta Ragoobeer, Tanguy Chambon, Bastiaan Cockx, Anna Júlia Éliás, Dóra Szabó, Máté Gugolya, Jan-Ulrich Kreft, Chloé Cloteau, Denisia Levercan, Mikaël Croyal, Thomas Delhaye, David Rondeau, Catherine Roullier, Samuel Bertrand, Carine Picot, Patrice Le Pape, Eric Batard, Nidia Alvarez-Rueda

**Affiliations:** Cibles et Médicaments des infections et de l’immunité, UR1155, IICiMed, Nantes Université.; Technical University of Denmark & University of Birmingham.; School of Biosciences & Institute of Microbiology and Infection, University of Birmingham, UK; Department of Morphology and Physiology, Faculty of Health Sciences, Semmelweis University, Hungary; Semmelweis University Doctoral School, Health Sciences Division, Semmelweis University, Hungary; Institute of Medical Microbiology, Faculty of Medicine, Semmelweis University, Hungary; HUN-REN-SU Human Microbiota Research Group, Budapest, Hungary; Nantes Université, CNRS, INSERM, l’institut du thorax, Nantes, France; Plateforme de Spectrométrie de Masse du CRNH-O, Nantes, France; Nantes Université, CHU Nantes, Inserm, CNRS, SFR Santé, Inserm UMS 016, CNRS UMS 3556, Nantes, France; Institut des Substances et Organismes de la Mer, ISOMer, UR 2160, Nantes Université, F-44000 Nantes, France; Nantes Université, Ecole Centrale Nantes, CNRS, LS2N, UMR 6004, F-44000 Nantes, France; Nantes Université, Université de Rennes, UMR CNRS 6164, IETR, F-35000 Rennes, France

**Keywords:** interkingdom interactions, *Candida* and *E. coli* interactions, multi-drug resistance

## Abstract

Interkingdom interactions between bacteria and fungi are an emerging research field that provides insights into pathological, environmental, and microbiota-related relationships. However, the mechanisms governing these interactions, particularly in the context of microbial resistance, remain largely unknown. This study aims to enhance our understanding of the complex interactions between different *Candida, Nakaseomyces and Sacharomyces* species from the human microbiota and two not isogenic strains of *Escherichia coli* (antibiotic-susceptible *E. coli*-ATCC and multidrug-resistant *E. coli*-OXA48). Forty-nine *Candida* strains were co-cultured with the two *E. coli* strains. Both bacterial and yeast growth was monitored using flow cytometry and compared to monocultures. The effect of *yeast* culture supernatants on *E. coli* proliferation was also investigated. Metabolomic fingerprints and metabolite identification were performed using mass spectrometry-based approaches followed by multiblock statistical analyses. The inhibitory powers (IP) of yeasts against *E. coli* and vice versa varied significantly among fungal species. *N. glabrata* exhibited the strongest inhibition against *E. coli*-ATCC, while *Candida lusitaniae, C. kefyr, C. krusei, C. tropicalis*, and *C. dubliniensis* showed lower IPs. *C. parapsilosis* and *Saccharomyces cerevisiae* had no inhibitory effects. Against *E. coli*-OXA48, most yeasts displayed no inhibition, except for *N. glabrata*. Conversely, *E. coli* inhibited yeast growth more effectively, particularly *Candida albicans*. Fungal supernatants from *S. cerevisiae, C. lusitaniae*, and *N. glabrata* showed the highest inhibitory effects on *E. coli*-ATCC, while *S. cerevisiae, C. krusei*, and *C. lusitaniae* were most effective against *E. coli*-OXA48. Unsupervised metabolite profiling data analysis with multiblock approach highlighted a clustering of samples according to yeast species. Regarding inhibitory power on *E. coli* (ATCC or OXA48), active supernatants tend to cluster together suggesting the presence of similar metabolites; some were further characterized. This study highlights the diverse interactions between *E. coli* and commensal yeasts. From an applied perspective, these findings pave the way for identifying probiotics or postbiotics with potential applications in combating multidrug-resistant bacteria through novel antimicrobial compounds.

## Introduction

The growing issue of antimicrobial resistance is of significant concern today, necessitating accelerated research into new therapeutics. The Gram-negative bacterium *Escherichia coli* poses a serious problem in human health (1). Multi-drug resistant (MDR) *E. coli* is included in the World Health Organization (WHO) list of priority antibiotic-resistant pathogens that represent a major threat to humans and for which new antibiotics and innovative therapeutic strategies are urgently needed (2). As a member of the *Enterobacteriaceae* family, it is responsible for numerous infections such as urinary tract infections, septicemia, neonatal meningitis, and intestinal infections, with increasing antibiotic resistance (3). Third and fourth-generation cephalosporins and carbapenem are commonly used antimicrobials to treat Gram-negative bacterial infections in humans. In *E. coli*, resistance to cephalosporins is typically mediated by extended-spectrum *β*-lactamases (ESBLs), which are encoded by plasmids easily transferred between bacterial species and capable of hydrolyzing these antibiotics. In addition to ESBLs, carbapenem resistance may also arise through enzymes like carbapenem-hydrolysing oxacillinases (OXA) or OXA48-like carbapenemases, which represent a threatening resistance mechanism. These enzymes are clinically concerning as they significantly limit therapeutic options. Intestinal carriage of ESBL *E. coli* is generally asymptomatic but persistent (4,5).

The gut microbiota serves as a reservoir for antimicrobial resistance genes (6). This microbiome is composed of billions of bacterial, viral, fungal, and parasitic micro-organisms, enabling both the production and the exchange of hundreds of small molecules (i.e. < 1,200 Da), commonly called metabolites (7). These microorganisms can be commensal, symbiotic, or even pathogenic (8). Studies on intestinal bacteria, or the intestinal microbiota, have received the most attention due to the abundance of bacterial flora in the intestine and their impact on individuals’ health. The composition of the microbiome and its variations influence human health and disease (9). Recent studies have shown that alterations in the microbiota are associated with obesity, depression, diabetes, and allergy (10). Limited attention has been paid to the fungal microbiome, or mycobiome, because fungi are often observed as a relatively small fraction of the gut microbial population (11). Fungi may be particularly underreported in some studies due to the limitations of 16S ribosomal RNA based methods (12). Recent studies used ITS gene regions for the sequencing of fungi in stool samples (13). Additionally, the analysis of the mycobiome is more challenging because many fungal species are difficult to culture in the laboratory and available analytical techniques are generally less developed compared to those available for bacteria (14). Fungal cells can be over 100 times larger than typical bacterial cells and are thus a more substantial part off the gut biomass than what sequencing based techniques would suggest (15), and they also play a role in health (11,12,16–19). Yeasts belonging to the genus *Candida* are among the most prevalent fungal genera within the microbiome, although the composition of the microbiome exhibits considerable variability between individuals (13). While *Candida* is generally non-pathogenic in healthy individuals, alterations in the microbiome combined with a compromised immune status can facilitate its colonization of tissues, leading to mucosal and dermal infections of varying severity. These infections can range from sepsis to those caused by biofilms on implanted medical devices (20). Similarly, alterations in the mycobiota are associated with certain pathologies. Specifically, research has demonstrated that patients with Crohn’s disease exhibit significantly elevated levels of the fungus *Candida tropicalis* compared to healthy family members (21).

There is currently little data on polymicrobial interactions within the human gut microbiome and their importance for health. To date, monomicrobial interactions (22) and polymicrobial interactions in the environment (23) have been studied. However, polymicrobial (bacterial-yeast) infections occur and increase the severity of infection or promote antimicrobial resistance (22,24). These cross-kingdom interactions exist within various niches of the human body, especially in the gut where they affect host health and diseases. Little is known about polymicrobial interactions in the gut, with *in vitro* data being predominant over *in vivo* data. These polymicrobial interactions play a role in host health. For instance, Sokol et al., (25) reported a significantly increased number of fungi–bacteria correlations in patients with ulcerative colitis compared to healthy individuals, suggesting an alteration in interkingdom crosstalk in this disease.

Given that pathophysiological interactions exist between *E. coli* and yeasts in the microbiota, a rational approach to characterising agonist and antagonist interactions is relevant in the context of combating microbial resistance. The main objective of this work was to develop an *in vitro* model for the characterisation of interactions between MDR *E. coli* and commensal yeasts belonging to different genera and species (*Candida*, *Saccharomyces,* and *Nakaseomyces*).

## Materials and methods

### Strains and culture conditions

The medium for growth curves and to count colonies was Sabouraud Chloramphenicol medium (Biokar diagnostics, France). The medium for yeast and bacterial cultures and co-cultures was Yeast Peptone Dextrose (YPD) medium (Sigma-Aldrich, USA). It was prepared from 10 g of yeast extract, 20 g of peptone, 900 mL of distilled water, and 100 mL of 10X dextrose (10g/L). It was then filtered using a stericup containing a 0.22 μm pore size filter to preserve as many nutrients as possible. Cefotaxime was diluted at 100 μg/mL in this medium for co-cultures with *E. coli*-OXA48 to maintain selection pressure.

Forty-nine commensal yeast strains from patients from Nantes University Hospital were used in these experiments: 19 strains of *Candida albicans* (Caal18, Caal21, Caal22, Caal*26,* Caal27, Caal31, Caal34, Caal38, Caal44, Caal45, Caal 48, Caal75, Caal86, Caal87, Caal93, Caal121, Caal123, Caal124, Caal146), 4 strains of *Nakaseomyces* (previously *Candida*) *glabrata* (Cagl1, Cagl2, Cagl5, Cagl7), 4 strains of *Candida tropicalis* (Catr1, Catr2, Catr3, Catr4), 4 strains of *Candida parapsilosis* (Capa2, Capa3, Capa5, Capa11), 4 strains of *Candida dubliniensis* (Cadu1, Cadu2, Cadu3, Cadu4), 4 strains of *Candida lusitaniae* (Calu1, Calu4, Calu5, Calu6), 4 strains of *Candida krusei* (Cakr2, Cakr4, Cakr5, Cakr7), 4 strains of *Candida kefyr* (Cake3, Cake4, Cake5, Cake6), 2 strains of *Saccharomyces cerevisiae* (Sace1, Sace2). They were stored at -80°C, then thawed on Sabouraud Chloramphenicol agar plates and stored at 4°C. Bacterial strains: An *E. coli* ATCC 25992 strain was used as antibiotic sensitive strain. The multi-drug resistance (MDR) *E. coli*-OXA48 strain isolated from a patient from Nantes University Hospital possess enzymes with hydrolytic activity against carbapenems and cephalosporins (C3G). Bacterial strains were stored at -80°C and then thawed on HBI agar or ESBL agar consisting of a nutrient base and an additional mixture of antibiotics, respectively.

### *In vitro* co-cultures between *E. coli* and yeasts

Yeasts were inoculated with a micropipette tip into 2 mL of YPD medium. After vortexing, the optical density at 600 nm was measured by taking 100 µL of each culture and checking that it reached an OD600 of approximately 0.1. The cultures were then incubated at 35°C until an OD600 of approximately 1 to reach exponential growth. Sensitive *E. coli* and resistant *E. coli*-OXA48 bacteria were inoculated using a micropipette tip into 2 mL of YPD and YPD + cefotaxime, respectively, and incubated under similar conditions than yeasts to reach exponential growth stage. The yeast precultures were then diluted in 5 mL of YPD medium and 5 mL of YPD + cefotaxime medium to obtain an OD600 = 0.05. The dilution was adapted to each strain. Bacterial pre-cultures were diluted to OD600 = 0.1 in 2 mL of YPD and YPD + cefotaxime. Bacteria were dispensed at 2 µL per well into flat-bottomed 96-well plates, and yeast was added at 200 µL per well, in addition to the yeast-only and bacteria-only controls. The UFC/mL of bacteria and yeasts is thus equal by these dilutions. At 0 h, 2 h, 4 h, 6 h, 8 h, and 10 h of co-culture, 200 µL of the corresponding time was transferred to a conical-bottom plate, centrifuged for 5 min at 2500 rpm and then were fixed with 200 µL of 4% PBS + paraformaldeyde. Finally, the cells were counted using a flow cytometer.

### Flow cytometry

Flow cytometry was used to measure changes in bacterial and yeast concentrations in pure and mixed cultures over time. The forward scatter (FSC) indicates the size of the particles (bacteria or yeast) in the sample. The FSC amplitude for yeasts was an order of magnitude higher than for the bacteria (for reference, the dynamic range of FSC values for the cytometer is approximately 6.5 orders of magnitude). Hence, with certain caveats, FSC can discriminate between the yeasts and bacteria used in this experiment. In addition to the FSC parameter, we used SYTO^®^ green-fluorescent nucleic acid stains to distinguish between live and dead cells in each population (bacteria and yeast) in mixed cultures. SYTO^®^ is a cell-permeable nucleic acid stain with enhanced fluorescence upon binding nucleic acids. SYTO^®^ was used to assess membrane integrity and cell viability, based on dye exclusion. The FSC *vs*. Side scatter (SSC) plots triggered the counting of individual particles (bacteria or yeast). Titers of *E. coli* and fungi were assessed by FACS until 10 h of co-culture. Titers of *E. coli* and fungi were assessed as the log10 of UFC/mL. The trapezoidal method was used to compute the area under the curve (AUC) of titers (of either *E. coli* or fungus, in log10 CFU/mL) against time. The inhibitory power of fungus against *E. coli* was defined as the ratio of the AUC of *E. coli* titers when *E. coli* was grown without fungus against the AUC of *E. coli* titers when *E. coli* was co-cultured with fungi (IP-Y/E). Inversely, the IP of *E. coli* against fungus was defined as the ratio of the AUC of fungus titers when fungus was grown alone against AUC of fungus titers when fungus was co-cultured with *E. coli* (IP-E/Y). Hence, growth inhibition was defined as an IP higher than 1.

### *In vitro* culture of *E. coli* in the presence of yeast culture supernatants

Bacteria and yeast were inoculated into 2 mL of YPD liquid medium and YPD + cefotaxime liquid medium. They were incubated at 35°C overnight. The OD600 was then measured and the yeasts were centrifuged at 4000 rpm for 10 min. The yeast supernatants were filtered, and the pH was measured for each supernatant to rule out the fact that a low pH, *i.e*. below 5, kills the bacteria. Then, in the wells of a 96-well plate, 2 μL of bacterial preculture was added to 200 μL of each supernatant. Each supernatant was made in triplicates, along with the bacteria in the YPD medium for the positive controls and the YPD medium alone for the negative control. OD600 was monitored for 10 h to obtain a growth curve. Each experiment was conducted on 3 different days, and each day, 3 wells were filled with one given culture supernatant and either *E. coli*-ATCC or *E. coli*-OXA48. The trapezoidal method was used to compute the AUC of the mean OD against time. The fungal supernatant IP (IP FS/E) was defined as the Log10 of the ratio of the AUC of *E. coli* titers when *E. coli* was grown without yeast supernatant against the AUC of *E. coli* titers when *E. coli* was grown with yeast supernatant. Unlike titers assessed by culture, OD could not be log-transformed due to zero values. Hence, the log-transformation was applied to the ratio of AUCs, and growth inhibition was defined by an IP_FS/E higher than 0.

### Metabolomic fingerprints of *Candida* supernatants by liquid chromatography-high-resolution mass spectrometry

All solvents were purchased from Biosolve (Valkenswaard, Netherlands). A quality control (QC) sample was prepared by pooling 50 μL from each individual sample and subsequently split into several aliquots. Metabolites were extracted from individual supernatant samples, QC samples and media (used as controls) by protein precipitation. Ice-cold methanol (800 µL) was added to defrost samples (200 µL). Following a 10-s vortex, the mixture was centrifuged (20,000 g, 10 min, 4°C), and the clear supernatant was split into two equal fractions of 300 µL. All samples were transferred to vials and then evaporated to dryness under a gentle nitrogen stream (room temperature). Dried samples were reconstituted either with 5% acetonitrile (fraction 1, 100 µL) or 75% acetonitrile (fraction 2, 100 µL), each containing 0.1% formic acid. The metabolome was analyzed by liquid chromatography-high-resolution mass spectrometry (LC-HRMS) on an Acquity H-Class^®^ UPLC^TM^ device (Waters Corporation, Milford, MA, USA) coupled to a Synapt^TM^ G2 HRMS Q-TOF mass spectrometer equipped with an electrospray ionization (ESI) interface, operating in both positive (ESI+) and negative (ESI-) ionization mode. Individual samples (5 μL) were randomized and injected altogether with QC samples either into a reversed-phased (RP) LC column (HSS T3, 2.1 × 100 mm, 1.7 µm, Waters Corporation; fraction 1) held at 50°C or a HILIC LC column (BEH-amide, 2.1 × 100 mm, 1.7 µm, Waters Corporation; fraction 2) held at 45°C. Metabolites from fraction 1 were separated with a linear gradient of mobile phase B (0.1% formic acid in acetonitrile) in mobile phase A (0.1% formic acid in water) at a flow rate of 400 μL/min. Mobile phase B was kept constant at 1% for 1 min, increased from 1% to 25% for 2.5 min, increased from 25% to 75% for 4.5 min, increased from 75% to 100% for 7 min, kept constant for 2 min, returned to the initial condition over 1 min, and kept constant for 5 min before the next injection. Metabolites from fraction 2 were separated with a linear gradient of mobile phase A (10 mmol/L ammonium acetate and 0.1% formic acid in water) in mobile phase B (98% acetonitrile, 10 mmol/L ammonium acetate, and 0.1% formic acid) at a flow rate of 400 μL/min. Mobile phase A was kept constant for 1 min at 1%, increased from 1% to 50% for 10 min, kept constant for 2 min, returned to the initial condition over 1 min, and kept constant for 4 min before the next injection. The full-HRMS mode was applied for metabolite detection (mass-to-charge ratio [*m/z*] range 200–1,200) at a mass resolution of 25,000 full-widths at half maximum (centroid mode). The ionization settings were as follows: capillary voltage, +3 kV (ESI+) or -2 kV (ESI-); cone voltage, 30 V; desolvation gas (N2) flow rate, 900 L/h; desolvation gas/source temperatures, 450°C / 120°C. Leucine enkephalin solution at 2 μg/mL (50% acetonitrile) was infused at a constant flow rate of 10 μL/min in the lock spray channel, allowing for correction of the measured *m/z* throughout the batch. Data acquisition and processing, including peak detection, integration, and alignment, were achieved using MassLynx^®^ and MakerLynx^®^ software, respectively (version 4.1, Waters Corporation). The relative standard deviation (RSD, %) was calculated for ion peak areas to highlight the repeatability of the analytical process in QCs. Variables (defined by a *m/z* ratio, a retention time, and an absolute ion peak area) having an RSD value below 30% on their absolute ion peak area were retained for further analysis (26).

### Multiblock analysis of metabolomic fingerprints of *Candida* supernatants

For metabolomics, the multivariate statistical analyses were performed using R 4.1.1 (CRAN https://cran.r-project.org/). The four data matrices (fraction 1 – RP – positive and negative ionization, and fraction 2 – HILIC positive and negative ionization) were analyzed simultaneously after high-level data fusion (27) by normalizing the total inertia of each block to 1 (28), providing a multiblock (MB) data matrix of 1,344 features detected in the 54 samples. Thus, unsupervised multiblock multivariate analysis was performed using ComDim (equivalent to PCA after multiblock correction) (28,29). Variable selection was performed using supervised multiblock partial least squares analysis (MB-PLS) using the package “ropls” (30) (multiblock correction, based on normalization of total block variance, was performed manually in R *prior* to multivariate analysis). The Y data for MB-PLS model construction were log-transformed and Unit Variance (UV) scaled IPs. Cross-comparison of MB-PLS models for variable selection was using the shared and unique structures (SUS) plot (31). Medians were reported with 1^st^ and 3^rd^ quartiles. The Kruskal-Wallis test was used to compare the medians of quantitative variables between groups. Correlations were assessed using the Spearman test. Differences were considered to be statistically significant at *P*-values < 0.05.

### Putative identification of metabolites from *Candida* supernatants based on metabolomic analyses

Metabolites were first identified and characterized by consulting the Human Metabolome Database (HMDB) (https://hmdb.ca/) and the Yeast Metabolome Database (YMDB) (http://www.ymdb.ca/) and then by the use of standard compounds (Sigma-Aldrich, Saint Quentin-Fallavier, France) with comparison to the exact mass measured, the elemental compositions with a mass error below 10 ppm (including specific adduct), the isotope patterns, the retention times according to the elution method, and the fragmentation patterns of selected precursor ion obtained by tandem mass spectrometry (MS/MS). According to these criteria, confidence levels were assigned to the putative identifications based on the Shymanski scale (32). For this study, only putatively identified metabolites with a confidence level of at least 4 were included.

### Statistical analysis

Medians were reported with inter-quartile ranges, and means with SD. Quantitative variables were compared using Wilcoxon test or Kruskal-Wallis test. Correlations were assessed using Spearman method. The inter- and intra-species variation was assessed using principal component analysis (PCA). A *P*-value < 0.05 was considered statistically significant. All statistical analyses were performed with R 4.2.3, R Foundation for Statistical Computing, Vienna, Austria.

## Results

### Growth curves variability of *Candida* and *E. coli* into cocultures

To investigate the interactions between the two *E. coli* strains and the 49 yeast strains belonging to the genera *Candida*, *Nakaseomyces*, and *Saccharomyces*, we conducted *in vitro* cultures in monoculture and co-culture in the YPD medium at 35°C without shaking for 10 h. Titers of *E. coli* and fungi were assessed as the log10 of UFC/mL. **Figure 1** depicts the log10 of UFC/mL over the 10 h of coculture incubation for each yeast strain against both *E. coli*-OXA48 and *E. coli*-ATCC strains. The results showed significant reduction between 4 and 10 h of fungi growth in coculture with *E. coli*-OXA48 and *E. coli*-ATCC strains (**Figure 1A-1B** and **Table S1**). In contrast, *E. coli*-OXA48 *E. coli*-ATCC growth curves significantly varied between 2 and 8 hours during coculture with yeast strains (**Table S2**). Interestingly, *E. coli*-OXA48 strain significantly overgrown between 2 and 4 h of coculture with yeast strains compared to monoculture controls and was then significantly inhibited between 6 and 8 h of coculture (**Figure 1C**). By contrast, growth of *E. coli*-ATCC strain was significantly reduced between 2 and 8 h of coculture compared to monoculture controls (**Figure 1D**).

**Figure 1.**
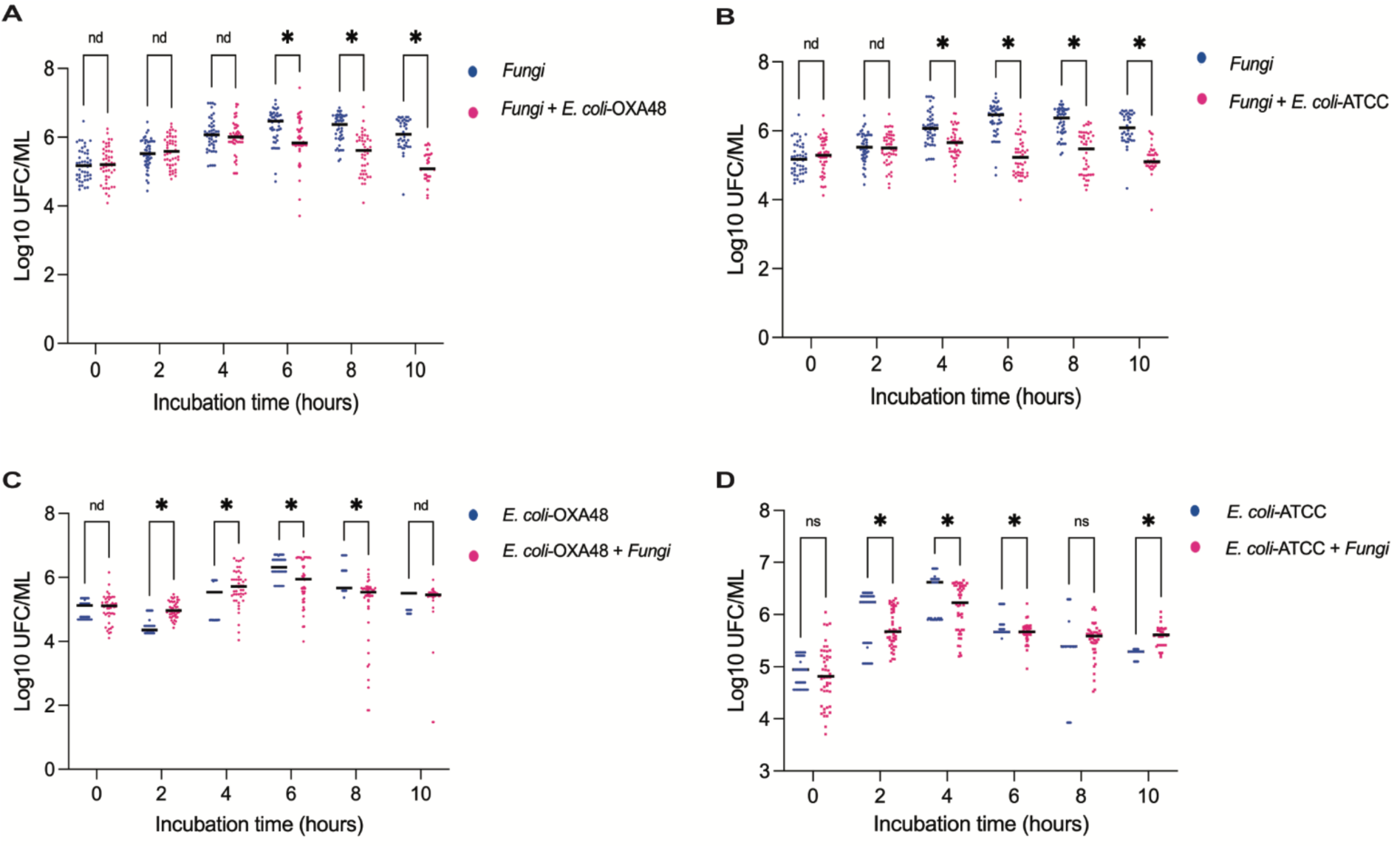
Dispersion of fungi and *E. coli* growth during incubation into cocultures. *E. coli*-ATCC, *E. coli*-OXA48 strains and the 49 yeast strains belonging to the genera *Candida*, *Nakaseomyces*, and *Saccharomyces*, were cultured *in vitro* in monoculture and co-culture in YPD medium at 35°C without shaking for 10 hours. The log10 of UFC/mL was calculated over the 10 h of coculture incubation for each *Candida* strain and both *E. coli*-OXA48 and *E. coli*-ATCC strains. Dots represent individual growth values from each coculture and experimental replicates from 49 yeast and *E. coli* strains. Bars represent median of the log10 of UFC/mL. Individual variance for each group was calculated and compared to respective control conditions (multiple unpaired t test with Welch correction, *P*-value < 0.05).

### Growth inhibition power (IP) of *Candida* and *E. coli* into cocultures

To further explain theses variations of growth under coculture conditions at a species and strain level, we defined a growth inhibition power (IP) for each bacterial and yeast strain into cocultures. The IP of the yeasts against *E. coli* was defined as the ratio of the AUCs of *E. coli* titers grown alone to *E. coli* titers in co-culture with yeast. Similarly, the IP of *E. coli* against fungi was calculated as the ratio of the AUCs of fungal titers grown alone to fungal titers in co-culture with *E. coli*. Growth inhibition was defined by an IP greater than 1 and under these conditions reflects either direct inhibition or competition for substrates, or both.

**Figure 2** depicts the inhibition variability between bacterial and fungal species. The IP values obtained for each fungal strain were different when co-cultured with *E. coli*-OXA48 and *E. coli*-ATCC, suggesting specific interactions at a species and strain level. Similarly, the two strains of *E. coli* exhibited significant IP variability across fungal species and strains. Although some yeast species and strains showed average IPs greater than 1, only the four strains of *N. glabrata* (Cagl) showed significant inhibition of *E. coli*-OXA48 with IPs ranging between 1.3 and 1.5 (**Figure 2A-2C**). Significantly, both *E. coli* strains displayed significant inhibitory power against all *C. albicans* strains, with IPs ranging between 1.1 and 1.4 (**Figure 2B**).

**Figure 2.**
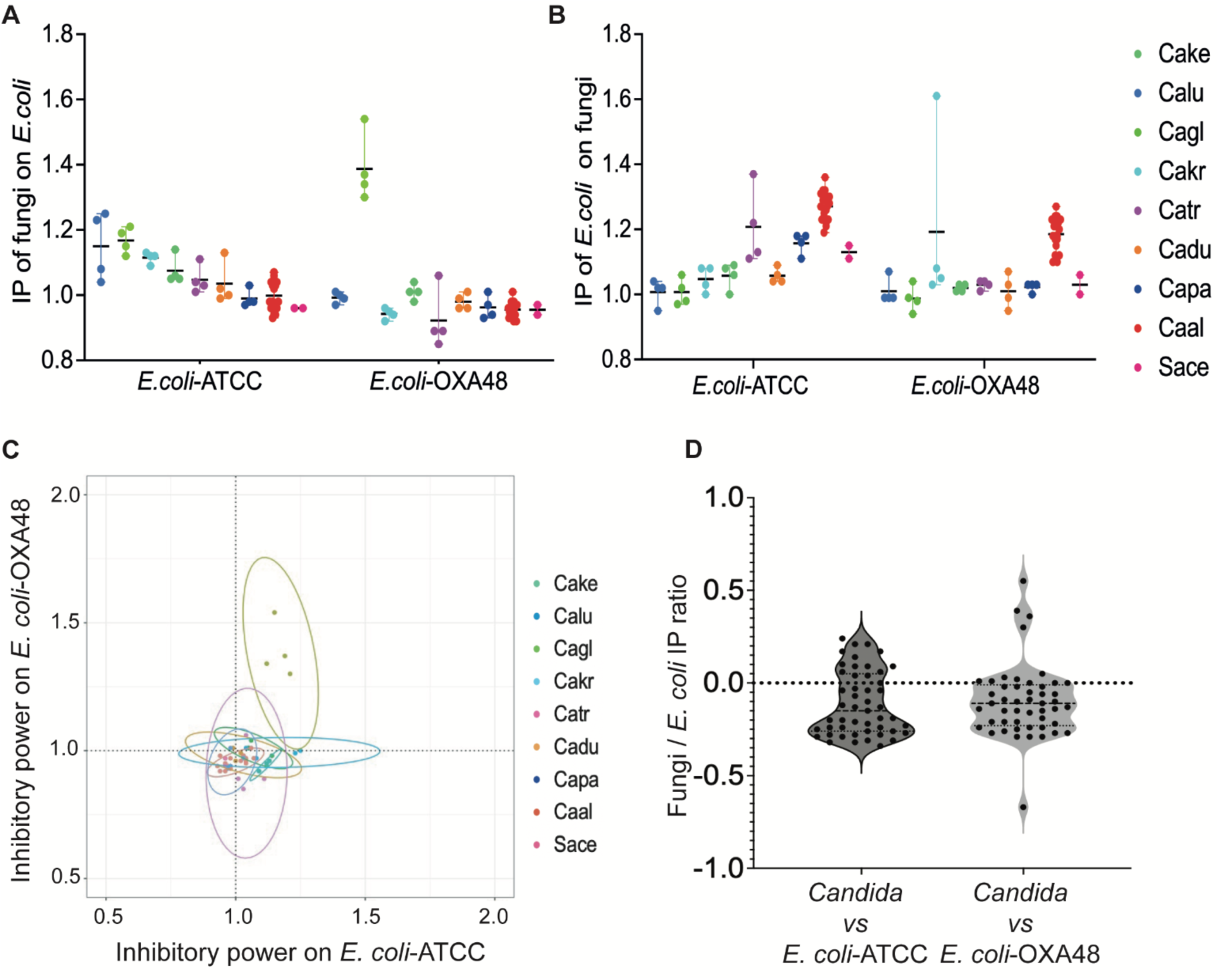
Inhibition power variability. *E. coli*-ATCC, *E. coli*-OXA48, and 49 yeast strains from genera *Candida*, *Nakaseomyces*, and *Saccharomyces* were cultured in YPD medium at 35°C for 10 hours without shaking. The inhibitory power (IP) of yeasts against *E. coli* was determined by the ratio of AUCs of *E. coli* grown alone to those grown with fungi. Similarly, the IP of *E. coli* against fungi was based on the AUCs of fungi grown alone to those with E. coli. An IP greater than 1 indicated growth inhibition. **(A)** IP of fungi on *E. coli*. **(B)** IP of *E. coli* on fungi. **(C)** Principal component analysis of inhibitory power of fungi against *E. coli* multiresistant and sensitive strains **(D)** The Fungi and *E. coli* ratio was calculated for each coculture. Dot colors represent the mean results of cocultures with each fungal strain from three independent experiments. The inter- and intra-species variation was assessed using principal component analysis (PCA). A *P*-value < 0.05 was considered statistically significant.

*In vitro*, the interactions between yeast and *E. coli* were antagonistic or neutral (**Figure 2D**). The IP ratios indicate that *E. coli* inhibits yeast strains more effectively. The mean IP of *E. coli* ATCC and *E. coli*-OXA48 against yeasts were highly correlated (Spearman rho: 0.71; *P*-value < 0.0001). The inhibition of *E. coli* by *Candida* varies depending on the species and strains. The mean inhibitory power of yeasts against *E. coli-*ATCC and *E. coli*-OXA48 were moderately correlated (Spearman rho: 0.47; *P*-value <0.001).

To define the interaction between *N. glabrata* and *E. coli* at the strain level, we found that the four *N. glabrata* strains grew equally well in the presence and in the absence of both *E. coli* strains (**Figure S1A-S1B**). Conversely, *E. coli*-OXA48 growth was specifically impacted by *N. glabrata*, showing growth inhibition by 2 h, suggesting direct inhibition (**Figure S1C**). Although there is a tendency for *N. glabrata* to inhibit the *E. coli*-ATCC strain, the results are not statistically significant (**Figure S1D**).

To define the interaction between *C. albicans* and *E. coli*, we found that the four *C. albicans* strains Caal18, Caal93, Caal121, were similarly inhibited by both *E. coli*-OXA48 and *E. coli*-ATCC strains (**Figure S2A-S2B**). The presence of *C. albicans* led to a faster apparent growth of *E. coli*-OXA48 between 0 and 4 h, while *E. coli*-ATCC growth rate seems to be equal during the first 4 h of incubation in the presence or in the absence of *C. albicans* (**Figure S2C-S2D** and **Table S3**).

### Inhibition of *E. coli* by fungal cultures supernatants

IP of supernatants (IP_FS/E) was defined as the log10 ratio of control AUC to culture supernatant AUC. An IP of 0 means both AUCs are equal, while an IP of 1 means the supernatant AUC is ten times lower than the control AUC. Median culture supernatant IP against *E. coli*-ATCC ranged from 0.03 (0.02-0.04) to 1.58 (1.48-2.51), exhibiting considerable variability (Kruskal-Wallis test *P*-value: 0.001; **Figure 3**). Similarly, IP against multidrug-resistant *E*. coli-OXA48 displayed significant variability (Kruskal-Wallis test *P*-value: 0.001), with values ranging from -0.03 (-0.05-0.02) to 1.62 (1.38-2.08). The three supernatants with the highest inhibitory effect on *E. coli*-ATCC were derived from *S. cerevisiae* (Sace2), *C. lusitaniae* (Calu5) and *N. glabrata* (Cagl5). For *E*. coli-OXA48, the most potent inhibitors were supernatants from *S. cerevisiae* (Sace2), *C. krusei* (Cakr7), and *C. lusitaniae* (Calu5).

**Figure 3.**
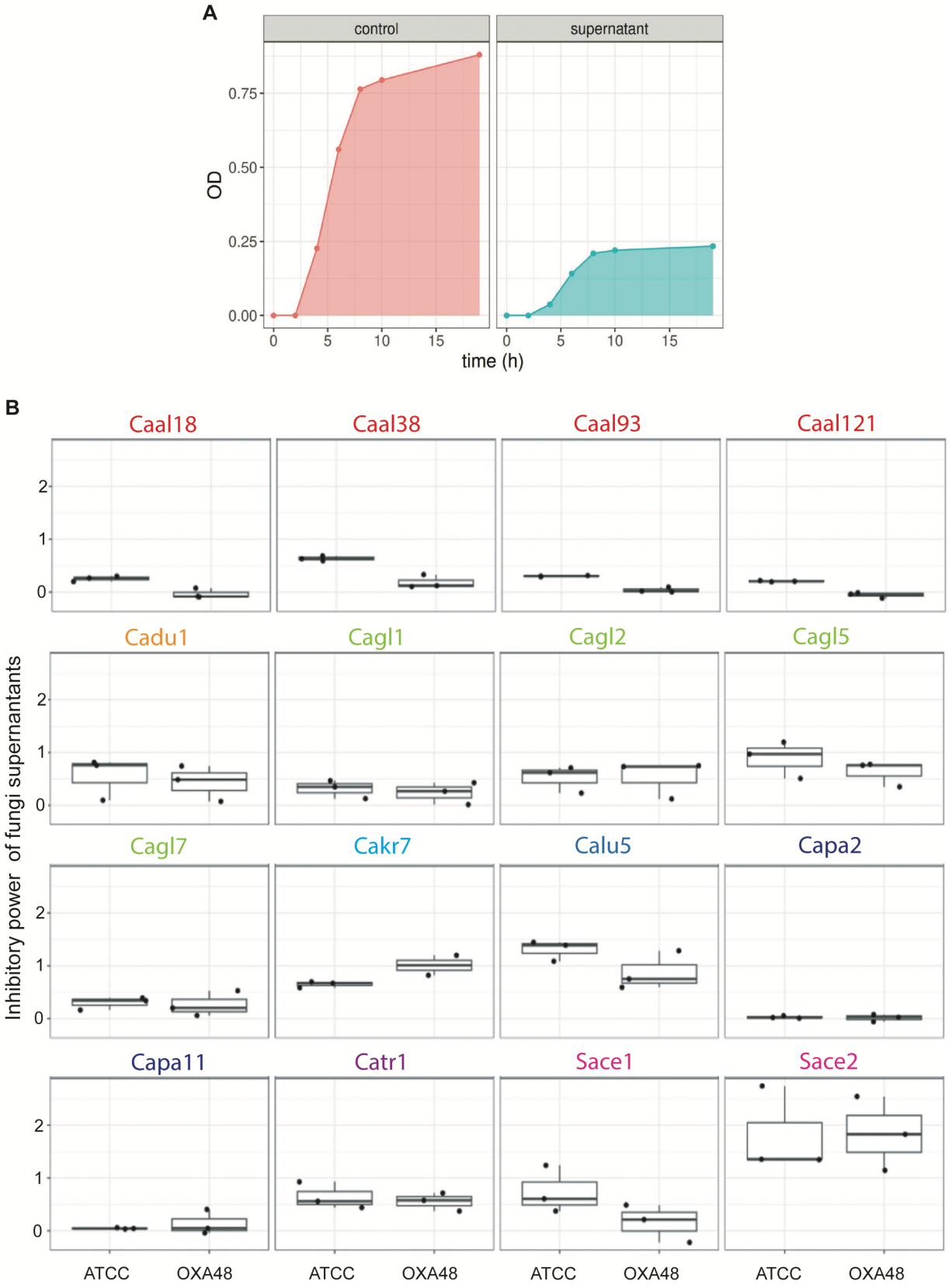
Inhibitory power of fungi supernatants against multi-resistant *E. coli-*ATCC and *E. coli-*OXA48. We selected 16 yeast strains that induced inhibition of *E. coli* growth in co-culture to test the effect of yeast culture supernatants on bacterial growth. **(A)** Representative growth curves of *E. coli* without fungal culture supernatant (control) and with culture supernatant. The area under the curve was computed for both conditions. The growth inhibition power (IP_FS/E) was defined as the AUCcontrol ratio against the AUCsupernatant. Fungal supernatants were obtained from three independent experiments. **(B)** An IP of 0 means both AUCs are equal, while an IP of 1 means the supernatant AUC is ten times lower than the control AUC. Each dot represents the mean results from three independent experiments, each in triplicate (Kruskal-Wallis test). A *P*-value < 0.05 was considered statistically significant. *C. albicans* (Caal18, Caal38, Caal93, Caal121); *C. dubliniensis* (Cadu1); *N. glabrata* (Cagl1, Cagl2, Cagl5, Cagl7); *C. krusei* (Cakr7); *C. lusitaniae* (Calu5); *C. tropicalis* (Catr1); *C. parapsilosis* (Capa2, Capa11); *S. cerevisiae* (Sace2).

Notable intra-species variability was observed for *Candida albicans* against *E. coli*-ATCC (Kruskal-Wallis test *P*-value < 0.05) and against *E*. *coli*-OXA48 (Kruskal-Wallis test *P*-value: 0.03), as well as for *S. cerevisiae* against *E*. coli-OXA48 (Kruskal-Wallis test *P*-value < 0.05).

Furthermore, the inhibitory efficacy against *E. coli*-ATCC and *E*. coli-OXA48 showed a high degree of correlation (Spearman rho: 0.82; *P*-value < 0.0001).

### IP correlation of fungi supernatant and IP of co-cultures against *E. coli*

Then we studied the correlation between the IPs of yeast supernatants and co-cultures against *E. coli* (**Figure 4**). There was no significant correlation between the 2 methods, either against *E. coli*-ATCC (Spearman rho: 0.25; *P*-value: 0.35) or against *E. coli*-OXA48 (Spearman rho: 0.34; *P*-value: 0.19).

**Figure 4.**
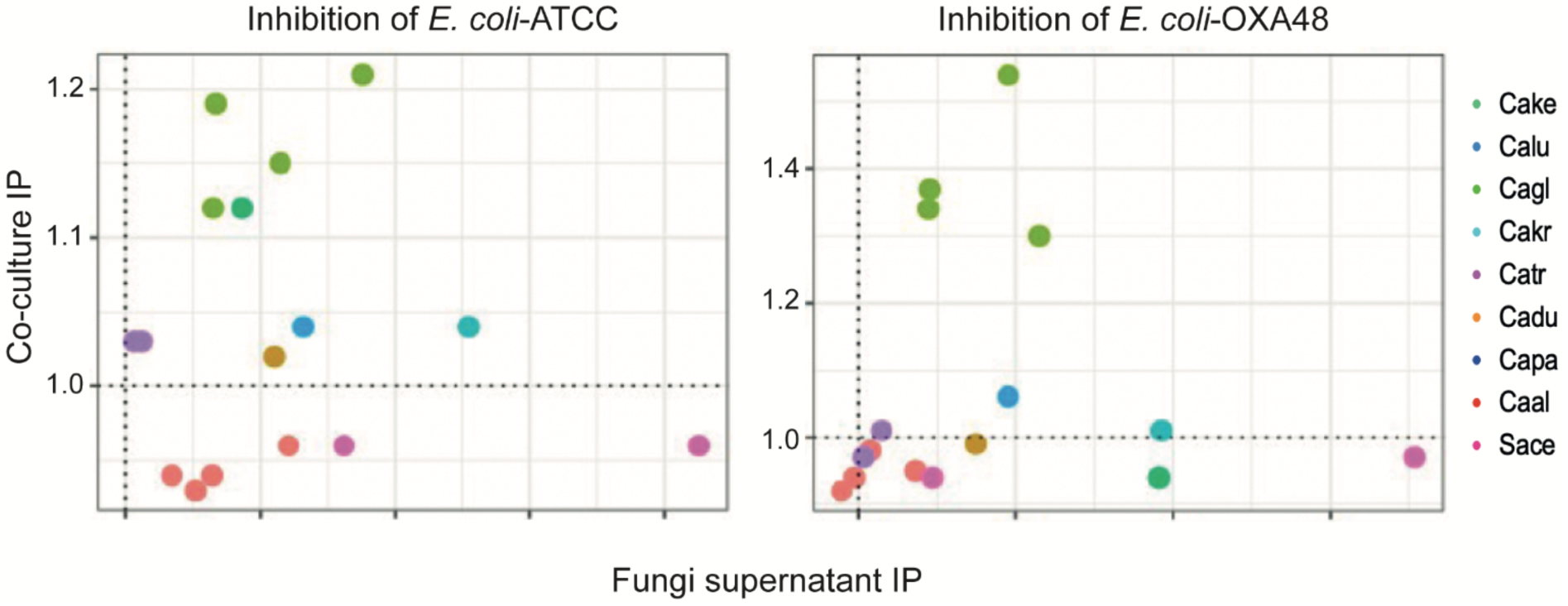
IP correlation of fungi supernatant and IP of co-cultures against *E. coli.* Yeast supernatants were incubated with the E. coli in YPD medium and YPD medium alone for the controls. OD600 was monitored for 10 h to obtain a growth curve. Each experiment was conducted on 3 different days, and each day, 3 wells were filled with one given culture supernatant and either *E. coli*-ATCC or *E. coli*-OXA48. The trapezoidal method was used to compute the area under the mean OD curve. The supernatant IP was defined as the log10-transformed ratio of control AUC against the AUC of the culture supernatant. Hence, higher IP was associated with lower ODs in the culture supernatant, and growth inhibition was defined by an IP higher than 0.

### Metabolomic analyses of fungal supernatants

To get an exhaustive overview of metabolic differences between supernatants, they were chemically profiled using LC-HRMS using four different methods (RP–ESI^+^, RP–ESI^-^, HILIC– ESI^+^, HILIC–ESI^-^). After data cleaning and filtration, the resulting data matrices were composed of 760, 315, 106 and 163 features for RP–ESI^+^, RP–ESI^-^, HILIC–ESI^+^, HILIC–ESI^-^ methods, respectively. To get an overall view of data organization, an unsupervised method merging the four sets of data was used. Thus, the ComDim (UV scaling) was used to highlight supernatants intrinsic similarities and/or difference. The ComDim score plot (**Figure S3**) provided an unsupervised visualization of the entire data structure. This analysis used intensity values from all peaks to account for nearly 80% of the sample variance, indicating good data representativeness. Samples from the same species grouped together, suggesting similar compositions. The first two components of the ComDim score plot accounted for 43.7 % and 23.1 % of the total variance (**Figure 5**). A preliminary clustering of the studied *Candida* species can be observed, even though the high number of species did not allow us to observe it significantly using only the first two dimensions. On this score plot, inhibition of the *E. coli-* ATCC and *E. coli-*OXA48 by fungal species was represented by dot size (**Figure 5A and 5B**, respectively) highlighting that inhibitory activity was not intrinsically related to data LC-HRMS organization. Other chemical signals could be related to such inhibition and should be found using a supervised multivariate approach.

**Figure 5.**
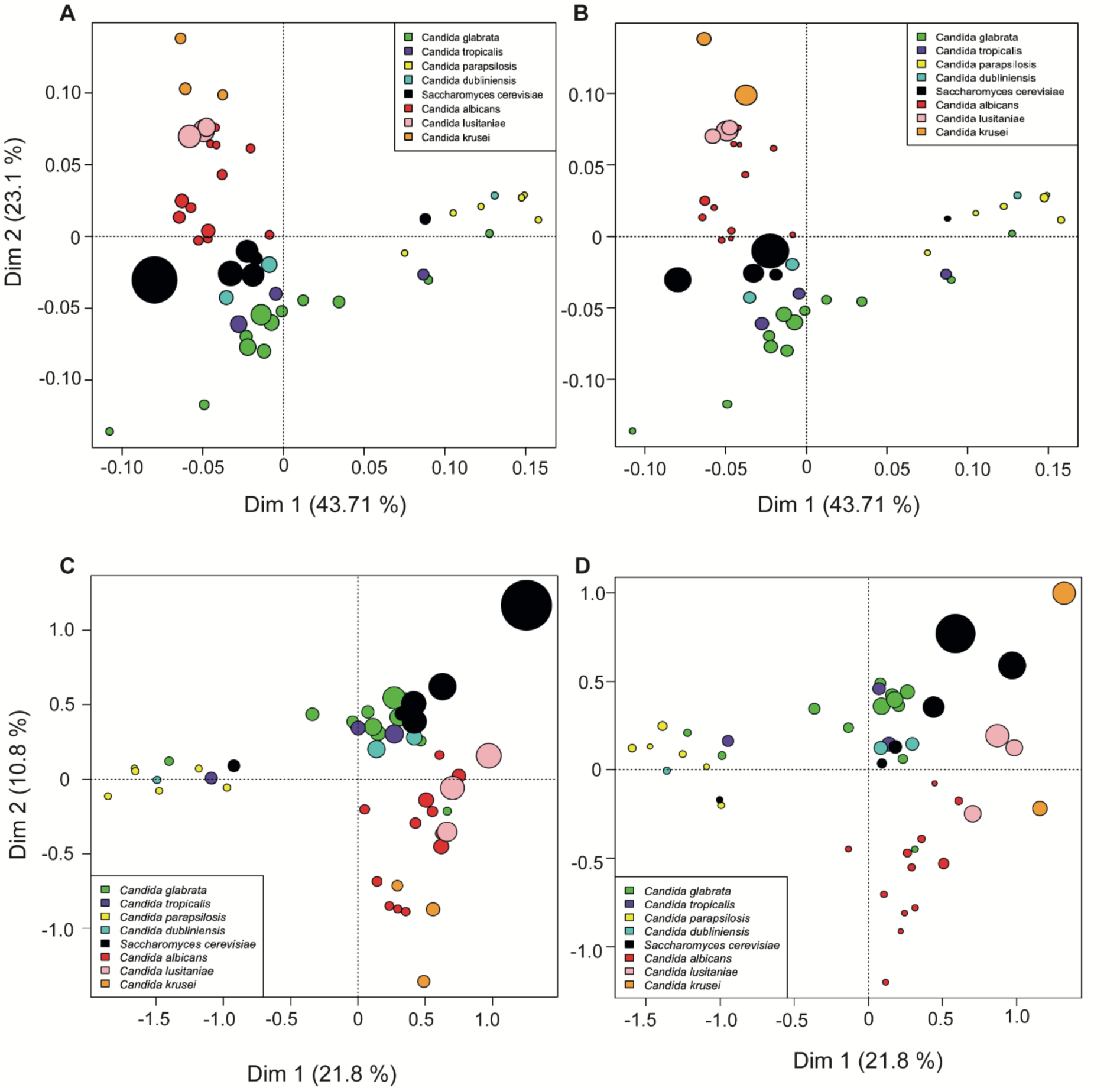
PCA score plot. LC-HRMS analyses (RP – ESI+, RP – ESI-, HILIC – ESI+, HILIC – ESI-) on fungal culture supernatants obtained from the 16 strains (Cagl1, Cagl2, Cagl5, Cagl7, Catr1, Capa2, Capa11, Cadu1, Caal18, Caal38, Caal93, Caal21, Calu5, Cakr7, Sace1, Sace2). Results correspond to three independent culture experiments, and color circles depict each fungal strain. Dot size is related to the activity of the fungal supernatants on the *E. coli* strain (as *log(PI)*). The largest dot corresponds to the most active supernatants. **(A)** ComDim score plot (dimensions 1 and 2) dot size was defined according to *log(PI)* against *E. coli*-ATCC. **(B)** ComDim score plot (dimensions 1 and 2) dot size was defined according to *log(PI)* against *E. coli*-OXA48. **(C)** MB-PLS score plot, modeling the *log(PI)*) against *E. coli*-ATCC. C) MB-PLS score plot, modeling the *log(PI)*) against *E. coli*-OXA48.

To highlight the features of interest from the data, two MB-PLS models (UV scaling) were constructed based on *Y* = log(*IP_E.coil-OXA48_*) and *Y* = log(*IP_E.coil-ATCC_*) (**Figure 5C**). Both models (*E. coli-*ATCC: model *P*-value 0.103 and *E. coli*-OXA48 model *P*-value 0.023) allowed to highlight features more present in supernatants having the highest biological activity (IP on the multi-drug-resistant *E. coli*-OXA48 and the susceptible *E. coli*-ATCC strains). The medium quality of the models reflects the species/strains specificity. Features of interest were selected based on Variable Importance in Projection (VIP) scores (> 2.5) and on their high presence within the active samples (MN scores). Finally, 9 features were significant for both *E. coli* strains, 6 exclusively for *E. coli*-ATCC, and 7 exclusively for *E. coli*-OXA48. Those 22 features were selected for further identification.

**Figure S2** depicts the Tyrosine identification by LC-HRMS to show the general strategy of metabolite identification. The HMDB analysis of the MS and MS/MS spectra used to identify and characterize peaks associated with significant inhibitory power on *E. coli*-ATCC and *E. coli*-OXA48. **Table 1** shows the metabolite annotation with a high level of confidence were presented (the complete list is presented in **Table S3**). As a result, 9 metabolites were distinctly identified, belonging to chemical families including fatty acyls, methylamines, amino acids, and acylglycerols. Five of them - oleate, stearate, palmitate, creatinine, and tyrosine were specific to *E. coli*-ATCC, while the remaining 4 - indole lactic acid, docosenamide, monoacylglycerol stearate [MG (18:0)], and glycerophosphocholine (GPC) - were common to both *E. coli*-ATCC and *E. coli*-OAX48. No metabolite specific to OAX48 was identified (**Table S3**).

**Table 1.** Proposed identification of metabolites of interest. MG: monoglycerol, GPC: glycerophosphocholine, CL: confidence level based on Shymanky scale, RT: retention time, STD: standard.

| ID | Adduct | Samples | Name | Family | C<br>L | R<br>T | M<br>S | MS<br>2 | ST<br>D |
| --- | --- | --- | --- | --- | --- | --- | --- | --- | --- |
| M1 | [M-H]<sup>-</sup> | ATCC | Oleate | Fatty acyl | 1 | ✓ | ✓ | ✓ | ✓ |
| M2 | [M-H]<sup>-</sup> | ATCC | Stearate | Fatty acyl | 1 | ✓ | ✓ | ✓ | ✓ |
| M3 | [M-H]<sup>-</sup> | ATCC | Palmitate | Fatty acyl | 1 | ✓ | ✓ | ✓ | ✓ |
| M4 | [M+H]<sup>+</sup> | ATCC | Creatinine | Methylamine | 1 | ✓ | ✓ | ✓ | ✓ |
| M5 | [M+H]<sup>+</sup> | ATCC | Tyrosine | Amino acid | 1 | ✓ | ✓ | ✓ | ✓ |
| M6 | [M+H]<sup>+</sup> | ATCC & OXA48 | Indole lactic acid | Indole | 1 | ✓ | ✓ | ✓ | ✓ |
| M7 | [M+H]<sup>+</sup> | ATCC & OXA48 | Docosenamide | Fatty amide | 2 | ✓ | ✓ | ✓ | X |
| M8 | [M+H]<sup>+</sup> | ATCC & OXA48 | MG (18:0) | Acylglycerol | 3 | ✓ | ✓ | ✓ | X |
| M9 | [M+H]<sup>+</sup> | ATCC & OXA48 | GPC | Methylamine | 4 | ✓ | ✓ | X | X |

MG : monoglycerol, GPC : glycerophosphocholine, CL : confidence level based on Shymanky scale, RT : retention time, MS, exact mass of precursor ion, MS/MS, fragmentation pattern in tandem mass spectrometry, STD : standard.

## Discussion

This study highlights the complex interactions between various yeast species and *E. coli* strains, including both antibiotic-susceptible (*E. coli*-ATCC) and multidrug-resistant (*E. coli*-OXA48) variants. *In vitro* co-culture experiments revealed that the IP of yeast against *E. coli*, and vice versa, varies significantly across species and strains. Notably, *N. glabrata* and *C. krusei* demonstrated the highest inhibitory effects against *E. coli*-ATCC, whereas most yeast strains showed limited or no inhibition against *E. coli*-OXA48, except for select strains of *N. glabrata* and *C. tropicalis*. Fungal supernatants exhibited diverse inhibitory effects on *E. coli* growth, with certain strains such as *S. cerevisiae*, *C. lusitaniae*, and *N. glabrata* showing potent activity. The high intra-species variability observed suggests that the inhibitory effects are strain-specific. Interestingly, while several fungal supernatants exhibited significant inhibitory effects on both *E. coli*-ATCC and *E. coli*-OXA48 strains, these effects were not consistently observed during direct co-culture experiments. This discrepancy suggests that secreted metabolites responsible for bacterial inhibition may accumulate to effective concentrations only under certain culture conditions or temporal dynamics not achieved in co-culture. In co-culture, competition for nutrients or rapid bacterial adaptation may reduce the efficacy of fungal-derived inhibitory compounds. Moreover, cell-cell contact and microenvironmental factors could modulate fungal metabolic pathways, potentially suppressing the production or activity of inhibitory metabolites during direct interaction. This hypothesis is supported by the lack of correlation between inhibition power (IP) measured from supernatants and from co-cultures, indicating distinct mechanisms at play. Similar findings have been described where microbial interactions altered bioactive molecule production, with metabolites only being effective in cell-free conditions (33). Further studies are needed to dissect the environmental or regulatory cues that enable or inhibit metabolite production during co-culture.

Interkingdom interactions between bacteria and fungi in the microbiome are an emerging area of interest because they can influence both the pathogenicity and antimicrobial resistance of these organisms. Our *in vitro* results are consistent with other studies that have shown that bacteria such as *E. coli* can interact with *Candida* species in ways that either enhance or inhibit their growth. Some studies have revealed a synergistic effect between bacteria and yeast. In particular, the fungal *ß*-1.3-glucan secreted increases the tolerance of *E. coli* to ofloxacin by preventing it from entering the biofilm (34). Studies indicate that the tolerance of *E. coli* to antimicrobials is facilitated by a small molecule of less than 10 kDa found in the supernatant of a *C. albicans* culture (24). Additionally, *E. coli* can enhance the virulent characteristics of *C. albicans* by increasing its biofilm formation capacity, its tolerance to nystatin, and the expression of its virulence genes *ALS3* and *HWP1*. Co-infection with *Galleria mellonella* larvae has been shown to increase mortality by up to 40% (35). Yang et al. (36) demonstrated that when *C. albic*ans and enterohaemorrhagic *E. coli* co-infected Caco-2 cells *in vitro*, *C. albicans* exhibited faster colonization, better accumulation, and up-regulation of its virulence genes compared to mono-infection, resulting in a more severe co-infection.

Other studies describe a rather antagonistic interaction between *E. coli* and *Candida*. Thein et al. (37) noted a significant negative correlation between the *in vitro* co-culture concentration of *E. coli* of oral origin and the number of viable yeasts in the biofilm, but only at high bacterial concentrations. Biofilm formation was inhibited under these conditions. Similarly, it was reported that *C. albicans* had lower biofilm formation when incubated with other types of bacteria, such as *E. coli*, than when incubated alone and this effect is the same when the bacteria are dead. The expression of hyphal-specific *C. albicans* genes was decreased (38). The interaction between *E. coli* and *C. albicans* also revealed a soluble fungicidal factor produced by *E. coli* that kills *C. albicans* in a magnesium-dependent manner and which could be a peptide (39). Finally, *E. coli* LPS is thought to have a direct modulating effect on the *in vitro* formation of biofilms of specific *Candida* species (40).

Exposure to an antibiotic or antifungal agent influences these interactions. It was shown that the colonization by *C. albicans* during a cefoperazone treatment was associated with a decrease in *Lactobacillus* populations and promoted the growth of *Bacteroidetes* during antibiotic recovery (41).

Mass spectrometry-based metabolomic analyses of fungal supernatants identified unique metabolic profiles linked to inhibitory activity. This study identified significant metabolites with potential roles in inhibition, such as oleate and indole lactic acid. Multiblock data analyses further refined these findings, isolating key metabolites with significant impacts on bacterial inhibition by capturing the diversity of peaks detected from the different types of LC-HRMS performed (reversed-phase and HILIC separation, positive and negative ionization modes). Supervised data analysis based on the IP of *E. coli* allowed the clustering of active supernatants of yeasts and highlighted the most important features explaining these differences. The differences observed could be attributed to their differential metabolism and point out genus and/or strain specificity within the response.

Fatty acids were associated with the inhibition of *E. coli* by fungal supernatants. Fatty acid biosynthesis in *Candida* species involves the production of both saturated and unsaturated fatty acids through the action of enzymes like fatty acid synthase (FAS). In fungi, the most common and abundant fatty acids (FAs) are palmitic acid (C16:0), stearic acid (C18:0), oleic acid (C18:1), and linoleic acid (C18:2). These can represent up to 95% of the total fatty acid content (42). Palmitate is one of the primary products of FAS in eukaryotic cells, including fungi. Stearate is typically synthesized during the elongation of fatty acid chains and serves as a precursor for other lipids, including unsaturated fatty acids such as oleate. The production of fatty acids, including oleate, is part of their normal lipid metabolism. This process can be influenced by various environmental factors such as substrate availability, culture conditions, and strain type. Certain *Candida* species, such as *C. albicans*, *C. tropicalis*, and *N. glabrata*, are also known for their ability to metabolize and produce lipids, which can include oleate. Oleic acid, plays several important roles in the microbiome, primarily by influencing the composition and function of microorganisms. Oleic acid has antimicrobial properties against certain pathogenic bacteria. It can interfere with the microbial cell membrane by altering its fluidity and disrupting membrane integrity, which prevents the growth of certain bacterial strains, including species of *Staphylococcus* and *Enterococcus* (43). Oleic acid can influence the composition of the gut microbiome by promoting the growth of beneficial bacteria, such as species of *Bifidobacterium* and *Lactobacillus*, which play an essential role in digestion and immune system regulation. On the other hand, it can also inhibit the growth of pathogenic bacteria, thus contributing to maintaining a healthy microbial balance in the intestinal tract. Oleic acid can affect the production of metabolites by microbiota microorganisms, including short-chain fatty acids (such as butyrate [C4:0]), which play a key role in gut health (44). Stearate and palmitate can be broken down or converted by specific gut bacteria into short-chain fatty acids (*e.g*., acetate [C2:0], propionate [C3:0], and butyrate [C4:0]). Monoacylglycerols are also lipid molecules that consist of a single fatty acid chain attached to a glycerol backbone and are intermediates in the lipid metabolism of many microorganisms, including yeasts.

Overall, this work provides a robust framework for understanding chemical and cell-cell interactions within yeast-*E. coli* and offers insights into potential bioactive metabolites that could be induced during such microbial co-culture (45). Understanding such chemical signaling could inform therapeutic strategies against multidrug-resistant bacteria.

---

**Figure S1.**
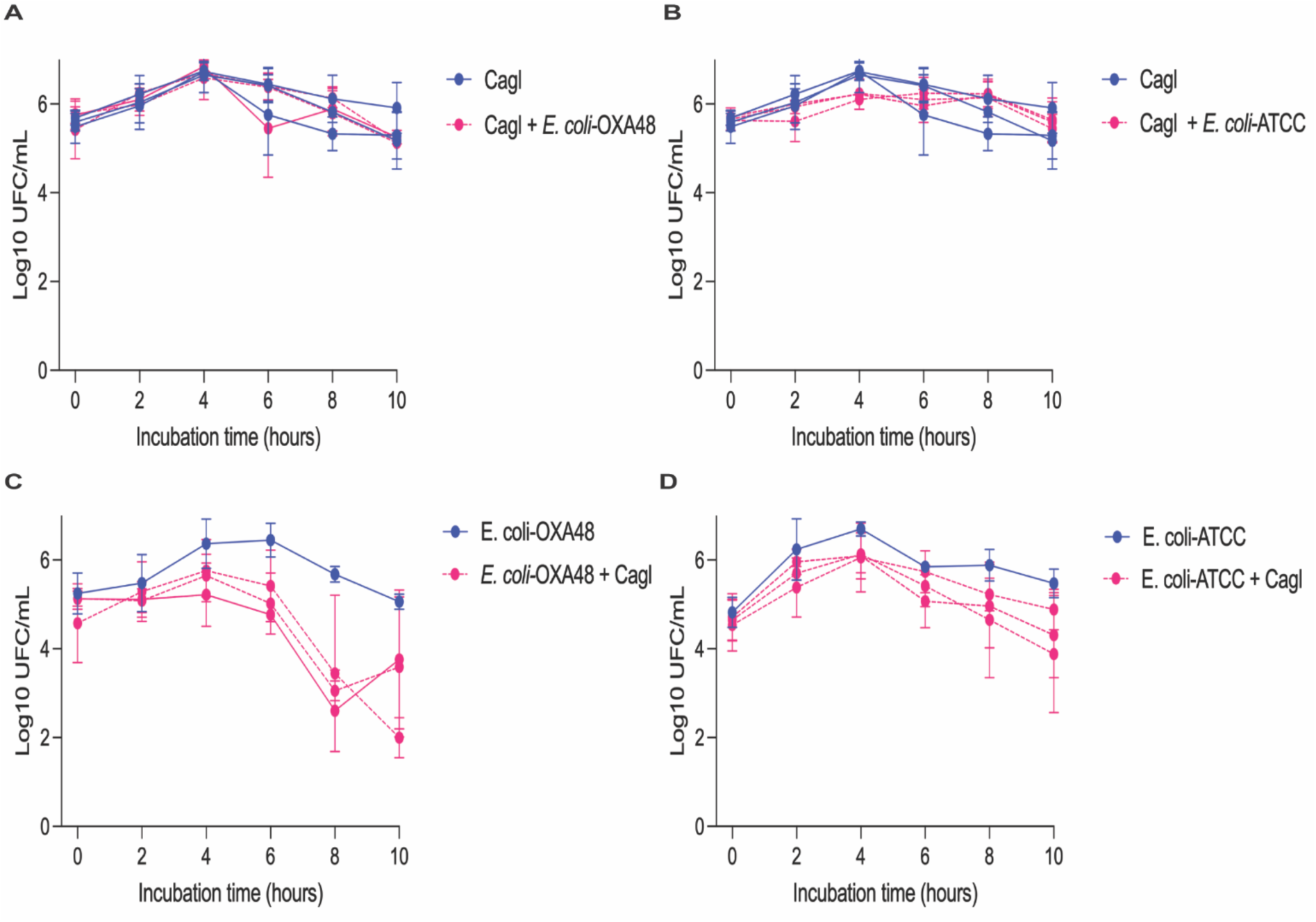
Growth inhibition of *E. coli*-ATCC and *E. coli*-OXA48 strains by *N. glabrata*. *E. coli*-ATCC, *E. coli*-OXA48 strains and four *N. glabrata* strains, were cultured *in vitro* in monoculture and co-culture in YPD medium at 35°C without shaking for 10 hours. The log10 of UFC/mL was calculated over the 10 h. Curves represent the microbial growth during coculture with each Cagl1, Cagl2, Cagl5 and Cagl7. Bars represent means ± SD of three independent experiments.

**Figure S2.**
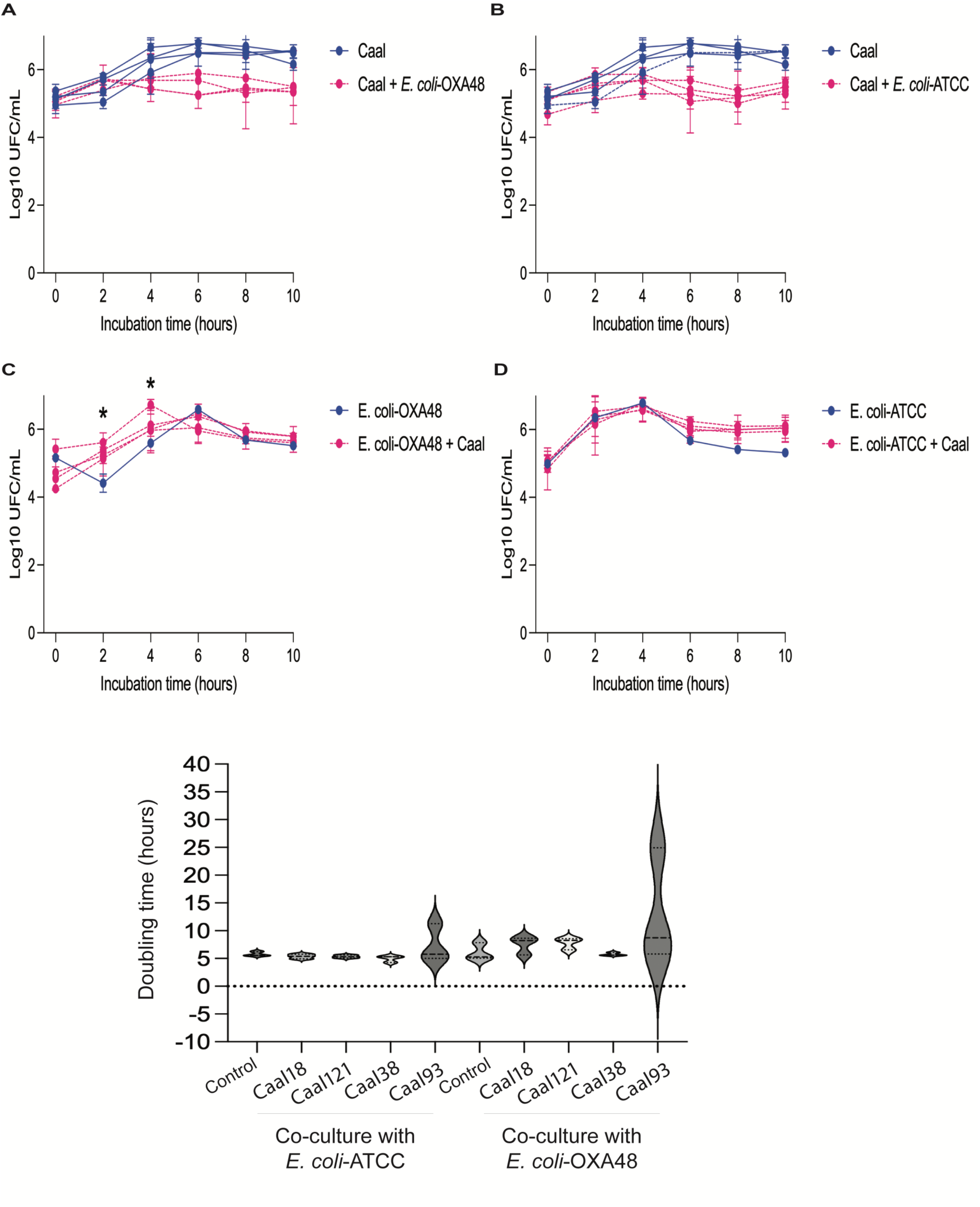
Growth inhibition of *C. albicans* by *E. coli*-ATCC and *E. coli*-OXA48 strains. *E. coli*-ATCC, *E. coli*-OXA48 strains and four *C. albicans* strains, were cultured *in vitro* in monoculture and co-culture in YPD medium at 35°C without shaking for 10 hours. The log10 of UFC/mL was calculated over the 10 h. Curves represent the microbial growth during coculture with each Caal18, Caal38, Caal93 and Caal121. Bars represent means ± SD of three independent experiments (2way ANOVA, *P*-value < 0.05).

**Figure S3.**
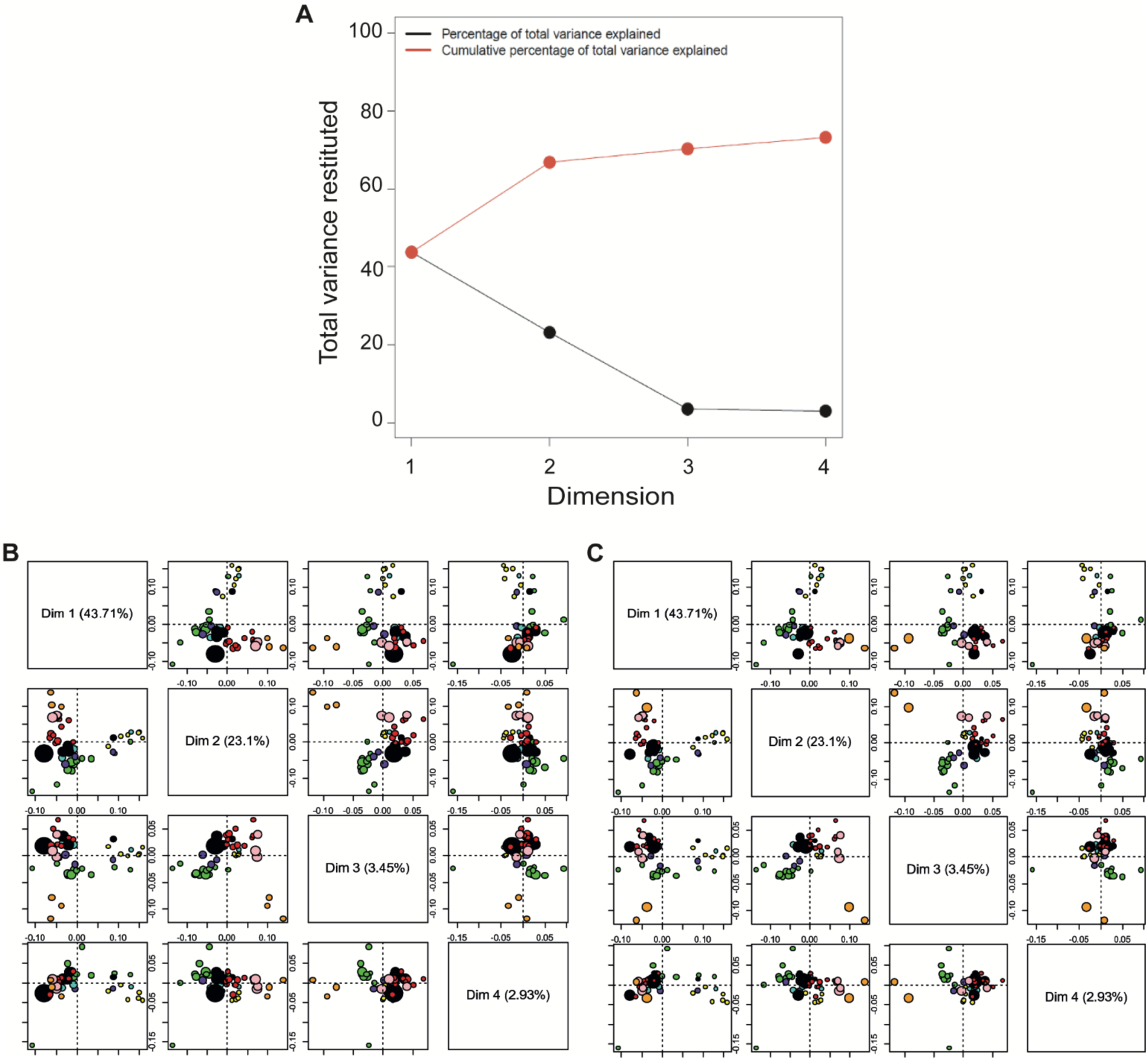
ComDim score plot (UV scaling). HPLC-MS analyses (RP – ESI+, RP – ESI-, HILIC – ESI+, HILIC – ESI-) on fungal culture supernatants obtained from the 16 strains (Cagl1, Cagl2, Cagl5, Cagl7, Catr1, Capa2, Capa11, Cadu1, Caal18, Caal38, Caal93, Caal21, Calu5, Cakr7, Sace1, Sace2). Results correspond to three independent culture experiments, and color circles depict each fungal strain. Circle size is related to the activity of the extracts on the *E. coli*-ATCC strain (as *log(PI)*) (the largest circle corresponds to most active extracts). **(A)** ComDim total explained variance for dimensions 1 to 4. **(B)** dot size related to *E. coli*-ATCC. **(C)** dot size related to multi-resistant *E. coli*-OXA48.

**Figure S4.**
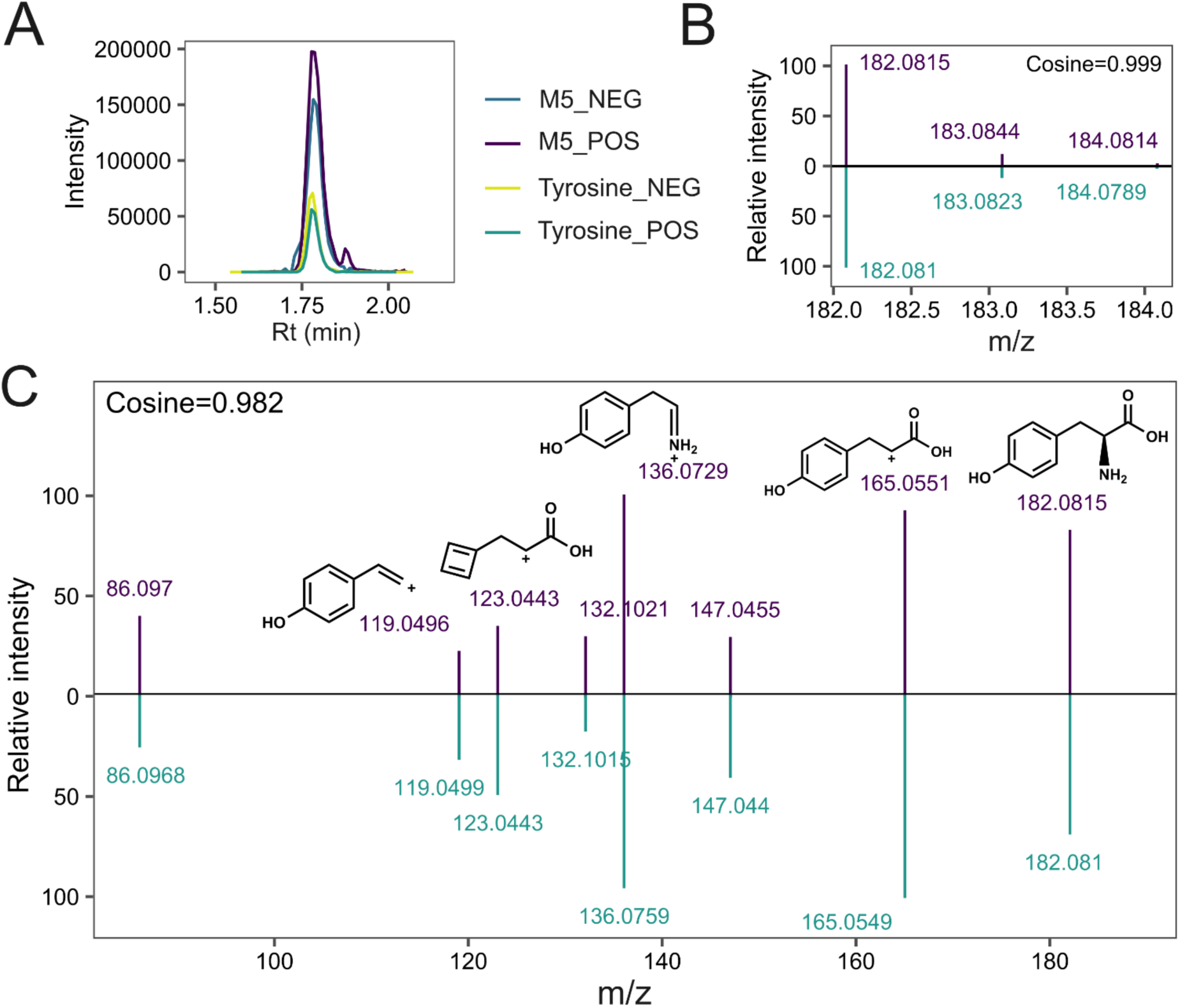
Tyrosine identification by LC-HRMS. **(A)** M5 and Tyrosine standard retention time after reversed-phase LC-HRMS (ESI+) and LC-HRMS (ESI-) acquisition. **(B)** MS^1^ spectra and fragmentation pattern of M5 (top) and Tyrosine standard (bottom) after LC-HRMS (ESI+) acquisition. **(C)** HMDB analysis of the MS/MS spectrum used to identify and characterize peaks associated with significant inhibitory power on *E. coli*-ATCC and *E. coli*-OXA48.

**Table S1.** Bacterial growth of *E. coli*-OXA48 and *E. coli*-ATCC over 10 h of incubation with overall fungi.

| Effect of Fungi on *E. coli*-OXA48 |  |  |  |  | Effect of fungi on *E. coli*-ATCC |  |  |  |
| --- | --- | --- | --- | --- | --- | --- | --- | --- |
| Incubation time (h) | Mean of *E. coli*-OXA48 | Mean of *E. coli*-OXA48 + fungi | Difference | *P* value | Mean of *E. coli*-ATCC | Mean of *E. coli*-ATCC+ fungi | Difference | *P* value |
| 0 | 4,998 | 5,028 | -0,02971 | 0,677488 | 4,903 | 4,832 | 0,07176 | 0,429476 |
| 2 | 4,459 | 4,987 | -0,528 | <0,000001 | 5,96 | 5,726 | 0,2343 | 0,013183 |
| 4 | 5,302 | 5,679 | -0,3765 | 0,000774 | 6,373 | 6,13 | 0,2434 | 0,007777 |
| 6 | 6,304 | 5,855 | 0,4483 | 0,000192 | 5,784 | 5,655 | 0,1289 | 0,001595 |
| 8 | 5,918 | 5,105 | 0,8124 | 0,000012 | 5,284 | 5,503 | -0,2189 | 0,058736 |
| 10 | 5,274 | 5,201 | 0,073 | 0,651942 | 5,265 | 5,581 | -0,3154 | <0,000001 |

Mean represent the log<sub>10</sub> of UFC/mL. Multiple unpaired t test with Welch correction, *P*-value < 0.05.

**Table S2.** Overall fungi growth over 10 h of incubation with *E. coli*-OXA48 and *E. coli*-ATCC strains.

| Effect of *E. coli*-OXA48 on fungi |  |  |  |  | Effect of *E. coli*-ATCC on fungi |  |  |  |
| --- | --- | --- | --- | --- | --- | --- | --- | --- |
| Incubation time (h) | Mean of fungi | Mean of fungi + *E. coli*-OXA48 | Difference | *P* value | Mean of fungi | Mean of fungi + *E. coli*-ATCC | Difference | *P* value |
| 0 | 5,19 | 5,197 | -0,006948 | 0,944735 | 5,19 | 5,284 | -0,09337 | 0,347628 |
| 2 | 5,529 | 5,579 | -0,04973 | 0,576946 | 5,529 | 5,482 | 0,04767 | 0,60694 |
| 4 | 6,074 | 6,002 | 0,0715 | 0,476549 | 6,074 | 5,62 | 0,4542 | 0,000006 |
| 6 | 6,305 | 5,888 | 0,4174 | 0,000638 | 6,305 | 5,232 | 1,073 | <0,000001 |
| 8 | 6,252 | 5,524 | 0,7286 | <0,000001 | 6,252 | 5,375 | 0,8769 | <0,000001 |
| 10 | 6,074 | 5,136 | 0,9378 | <0,000001 | 6,074 | 5,186 | 0,8881 | <0,000001 |

Means represent the log<sub>10</sub> of UFC/mL. Multiple unpaired t test with Welch correction, *P*-value < 0.05.

**Table S3.** Growth of yeast (*Candida*, *Nakaseomyces*, and *Saccharomyces* strains) and bacteria (with *E. coli*-OXA48 and *E. coli*-ATCC strains) over 10 h of co-culture incubation.

| Fungi strain | Effect of Fungi on *E. coli*-OXA48 |  |  | Effect of Fungi on *E. coli*-ATCC |  |  | Effect of *E. coli*-OXA48 on Fungi |  |  | Effect of *E. coli*-ATCC on Fungi |  |  |
| --- | --- | --- | --- | --- | --- | --- | --- | --- | --- | --- | --- | --- |
|  | Control median | Coculture median | *P* value | Control median | Coculture median | *P* value | Control median | Coculture median | *P* value | Control median | Coculture median | *P* value |
| Cagl1 | 5.58 | 5.05 | **0.0411** | 5.86 | 5.30 | 0.1797 | 6.04 | 5.89 | 0.5887 | 6.04 | 5.87 | 0.5887 |
| Cagl2 | 5.58 | 4.94 | **0.0152** | 5.86 | 5.02 | 0.2403 | 5.72 | 5.82 | 0.9372 | 5.72 | 5.98 | 0.6991 |
| Cagl5 | 5.58 | 4.94 | **0.0152** | 5.86 | 5.02 | 0.2403 | 5.72 | 5.82 | 0.9372 | 5.72 | 5.98 | 0.6991 |
| Cagl7 | 5.58 | 4.93 | 0.132 | 5.86 | 5.03 | 0.0931 | 5.93 | 6.19 | 0.8182 | 5.93 | 6.02 | >0.9999 |
| Catr1 | 5.58 | 5.75 | 0.6991 | 5.86 | 5.69 | 0.5887 | 6.01 | 5.83 | 0.8182 | 6.01 | 5.41 | 0.2403 |
| Capa2 | 5.58 | 5.73 | 0.8182 | 5.78 | 5.79 | 0.8182 | 5.74 | 5.57 | 0.8182 | 5.74 | 5.22 | **0.0411** |
| Capa11 | 5.58 | 5.65 | >0.9999 | 5.78 | 5.88 | >0.9999 | 5.71 | 5.69 | >0.9999 | 5.71 | 5.24 | **0.0152** |
| Dadu1 | 5.58 | 5.73 | 0.9372 | 5.78 | 5.71 | 0.8182 | 6.16 | 6 | 0.9372 | 6.16 | 5.76 | 0.3095 |
| Caal18 | 5.55 | 5.87 | 0.4848 | 5.54 | 6.18 | 0.4848 | 6.25 | 5.38 | **0.0411** | 6.25 | 5.57 | 0.0649 |
| Caal38 | 5.54 | 5.85 | 0.5887 | 5.52 | 5.98 | 0.5887 | 6.21 | 5.63 | 0.3095 | 6.21 | 5.18 | 0.1797 |
| Caal93 | 5.33 | 5.7 | 0.8182 | 5.56 | 6.08 | 0.2403 | 6.58 | 5.39 | **0.0152** | 6.58 | 5.37 | **0.0411** |
| Caal121 | 5.33 | 5.65 | 0.2403 | 5.56 | 6.02 | 0.1797 | 6.36 | 5.58 | 0.1797 | 6.36 | 5.33 | 0.0649 |
| Calu5 | 5.54 | 5.48 | 0.9372 | 5.52 | 5.73 | 0.9372 | 5.88 | 6.02 | 0.9372 | 5.88 | 6.17 | 0.1797 |
| Cakr7 | 5.54 | 5.78 | 0.3095 | 5.53 | 5.31 | 0.3095 | 6.38 | 4.79 | **0.0022** | 6.38 | 5.93 | 0.2403 |
| Sace1 | 5.53 | 5.82 | 0.5887 | 5.52 | 5.96 | 0.4848 | 6.02 | 6.07 | 0.8182 | 6.03 | 5.45 | 0.0649 |
| Sace2 | 5.55 | 5.77 | 0.3939 | 5.54 | 5.96 | 0.4848 | 6.22 | 5.96 | 0.3939 | 6.22 | 5.78 | 0.0649 |

Means represent all the log<sub>10</sub> of UFC/mL during 10 h of coculture incubation. Unpaired t test with Welch correction, *P*-value < 0.05.

**Table S4.**
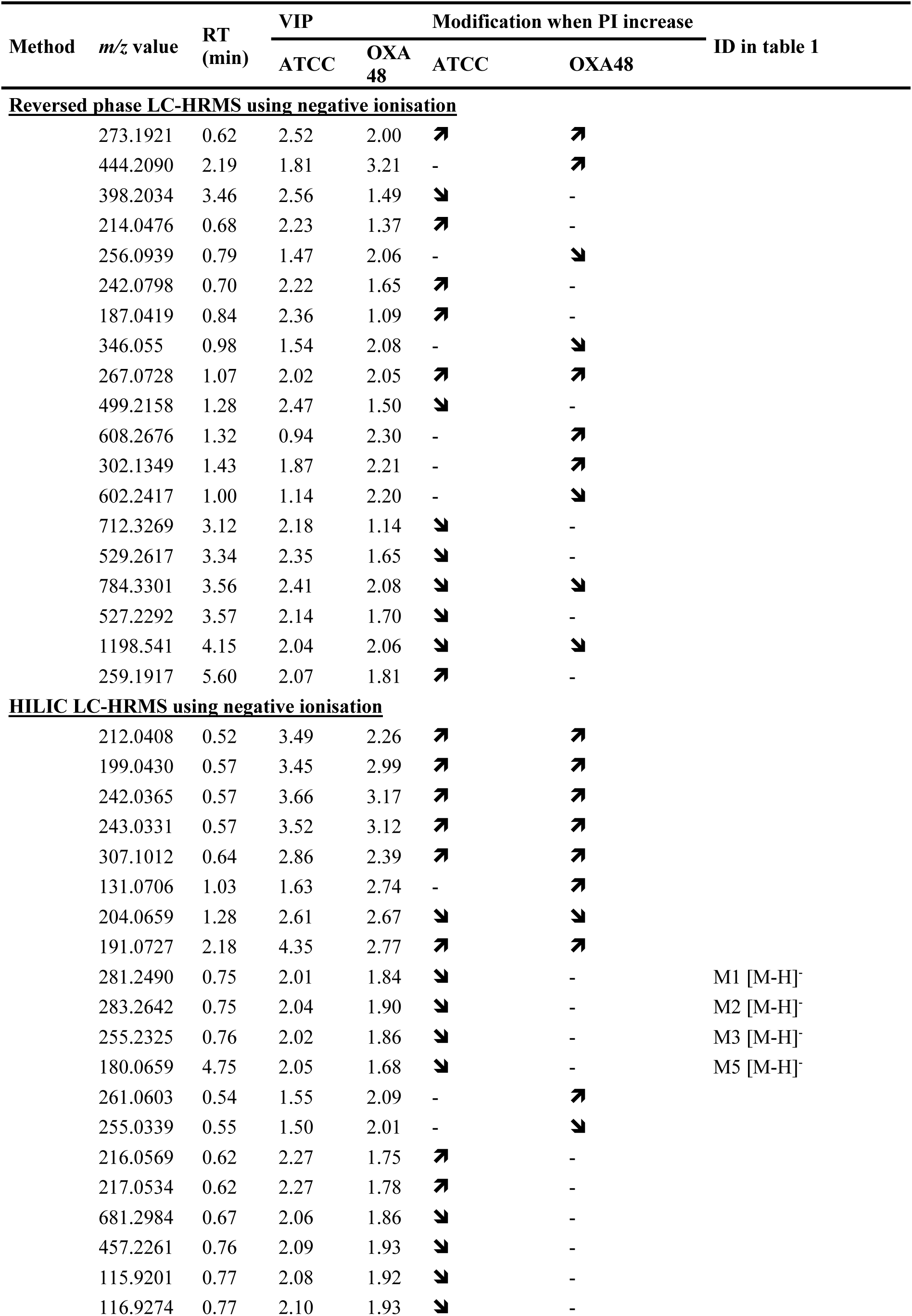
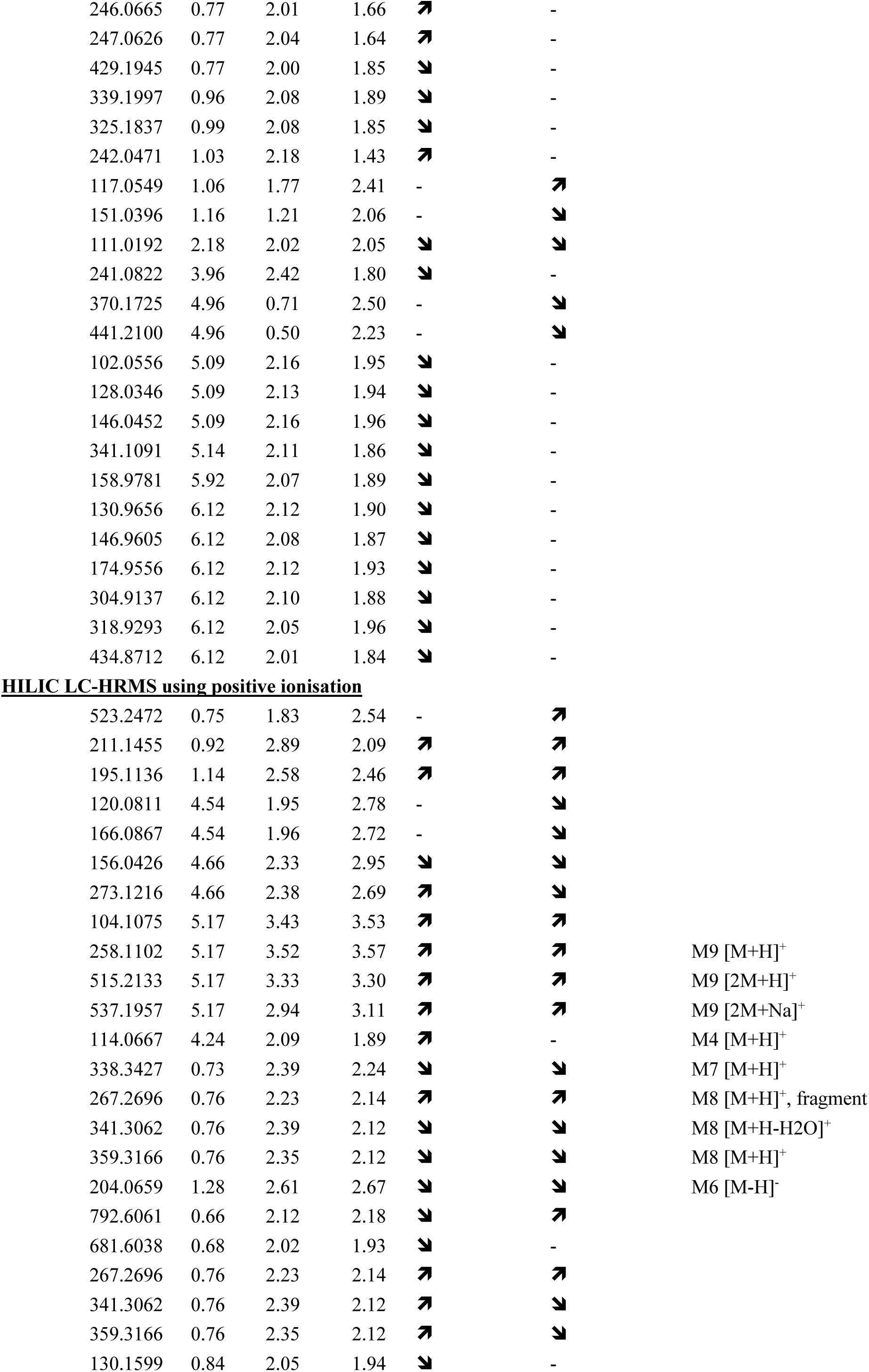
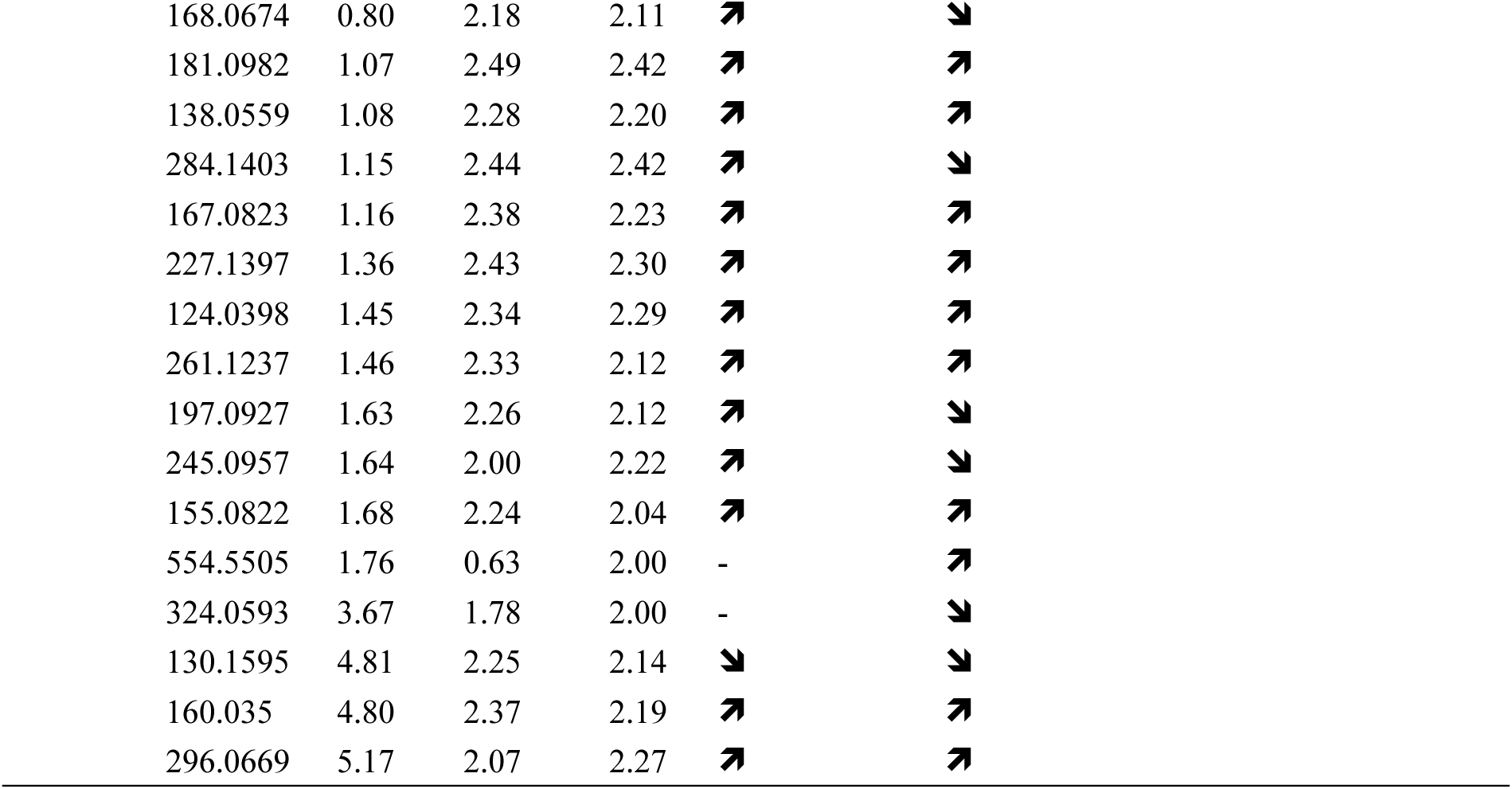
Features of interest based on the MB-PLS models using 640 VIP < 2.5. RT: Retention time.

| Method | *m/z* value | RT<br>(min) | VIP |  | Modification when PI increase |  | ID in table 1 |
| --- | --- | --- | --- | --- | --- | --- | --- |
|  |  |  | ATCC | OXA<br>48 | ATCC | OXA48 |  |
| Reversed phase LC-HRMS using negative ionisation |  |  |  |  |  |  |  |
|  | 273.1921 | 0.62 | 2.52 | 2.00 | ↗ | ↗ |  |
|  | 444.2090 | 2.19 | 1.81 | 3.21 | - | ↗ |  |
|  | 398.2034 | 3.46 | 2.56 | 1.49 | ↘ | - |  |
|  | 214.0476 | 0.68 | 2.23 | 1.37 | ↗ | - |  |
|  | 256.0939 | 0.79 | 1.47 | 2.06 | - | ↘ |  |
|  | 242.0798 | 0.70 | 2.22 | 1.65 | ↗ | - |  |
|  | 187.0419 | 0.84 | 2.36 | 1.09 | ↗ | - |  |
|  | 346.055 | 0.98 | 1.54 | 2.08 | - | ↘ |  |
|  | 267.0728 | 1.07 | 2.02 | 2.05 | ↗ | ↗ |  |
|  | 499.2158 | 1.28 | 2.47 | 1.50 | ↘ | - |  |
|  | 608.2676 | 1.32 | 0.94 | 2.30 | - | ↗ |  |
|  | 302.1349 | 1.43 | 1.87 | 2.21 | - | ↗ |  |
|  | 602.2417 | 1.00 | 1.14 | 2.20 | - | ↘ |  |
|  | 712.3269 | 3.12 | 2.18 | 1.14 | ↘ | - |  |
|  | 529.2617 | 3.34 | 2.35 | 1.65 | ↘ | - |  |
|  | 784.3301 | 3.56 | 2.41 | 2.08 | ↘ | ↘ |  |
|  | 527.2292 | 3.57 | 2.14 | 1.70 | ↘ | - |  |
|  | 1198.541 | 4.15 | 2.04 | 2.06 | ↘ | ↘ |  |
|  | 259.1917 | 5.60 | 2.07 | 1.81 | ↗ | - |  |
| HILIC LC-HRMS using negative ionisation |  |  |  |  |  |  |  |
|  | 212.0408 | 0.52 | 3.49 | 2.26 | ↗ | ↗ |  |
|  | 199.0430 | 0.57 | 3.45 | 2.99 | ↗ | ↗ |  |
|  | 242.0365 | 0.57 | 3.66 | 3.17 | ↗ | ↗ |  |
|  | 243.0331 | 0.57 | 3.52 | 3.12 | ↗ | ↗ |  |
|  | 307.1012 | 0.64 | 2.86 | 2.39 | ↗ | ↗ |  |
|  | 131.0706 | 1.03 | 1.63 | 2.74 | - | ↗ |  |
|  | 204.0659 | 1.28 | 2.61 | 2.67 | ↘ | ↘ |  |
|  | 191.0727 | 2.18 | 4.35 | 2.77 | ↗ | ↗ |  |
|  | 281.2490 | 0.75 | 2.01 | 1.84 | ↘ | - | M1 [M-H]<sup>-</sup> |
|  | 283.2642 | 0.75 | 2.04 | 1.90 | ↘ | - | M2 [M-H]<sup>-</sup> |
|  | 255.2325 | 0.76 | 2.02 | 1.86 | ↘ | - | M3 [M-H]<sup>-</sup> |
|  | 180.0659 | 4.75 | 2.05 | 1.68 | ↘ | - | M5 [M-H]<sup>-</sup> |
|  | 261.0603 | 0.54 | 1.55 | 2.09 | - | ↗ |  |
|  | 255.0339 | 0.55 | 1.50 | 2.01 | - | ↘ |  |
|  | 216.0569 | 0.62 | 2.27 | 1.75 | ↗ | - |  |
|  | 217.0534 | 0.62 | 2.27 | 1.78 | ↗ | - |  |
|  | 681.2984 | 0.67 | 2.06 | 1.86 | ↘ | - |  |
|  | 457.2261 | 0.76 | 2.09 | 1.93 | ↘ | - |  |
|  | 115.9201 | 0.77 | 2.08 | 1.92 | ↘ | - |  |
|  | 116.9274 | 0.77 | 2.10 | 1.93 | ↘ | - |  |

|  |  |  |  |  |  |
| --- | --- | --- | --- | --- | --- |
| 246.0665 | 0.77 | 2.01 | 1.66 | ↗ | - |
| 247.0626 | 0.77 | 2.04 | 1.64 | ↗ | - |
| 429.1945 | 0.77 | 2.00 | 1.85 | ↘ | - |
| 339.1997 | 0.96 | 2.08 | 1.89 | ↘ | - |
| 325.1837 | 0.99 | 2.08 | 1.85 | ↘ | - |
| 242.0471 | 1.03 | 2.18 | 1.43 | ↗ | - |
| 117.0549 | 1.06 | 1.77 | 2.41 | - | ↗ |
| 151.0396 | 1.16 | 1.21 | 2.06 | - | ↘ |
| 111.0192 | 2.18 | 2.02 | 2.05 | ↘ | ↘ |
| 241.0822 | 3.96 | 2.42 | 1.80 | ↘ | - |
| 370.1725 | 4.96 | 0.71 | 2.50 | - | ↘ |
| 441.2100 | 4.96 | 0.50 | 2.23 | - | ↘ |
| 102.0556 | 5.09 | 2.16 | 1.95 | ↘ | - |
| 128.0346 | 5.09 | 2.13 | 1.94 | ↘ | - |
| 146.0452 | 5.09 | 2.16 | 1.96 | ↘ | - |
| 341.1091 | 5.14 | 2.11 | 1.86 | ↘ | - |
| 158.9781 | 5.92 | 2.07 | 1.89 | ↘ | - |
| 130.9656 | 6.12 | 2.12 | 1.90 | ↘ | - |
| 146.9605 | 6.12 | 2.08 | 1.87 | ↘ | - |
| 174.9556 | 6.12 | 2.12 | 1.93 | ↘ | - |
| 304.9137 | 6.12 | 2.10 | 1.88 | ↘ | - |
| 318.9293 | 6.12 | 2.05 | 1.96 | ↘ | - |
| 434.8712 | 6.12 | 2.01 | 1.84 | ↘ | - |

**HILIC LC-HRMS using positive ionisation**
|  |  |  |  |  |  |  |
| --- | --- | --- | --- | --- | --- | --- |
| 523.2472 | 0.75 | 1.83 | 2.54 | - | ↗ |  |
| 211.1455 | 0.92 | 2.89 | 2.09 | ↗ | ↗ |  |
| 195.1136 | 1.14 | 2.58 | 2.46 | ↗ | ↗ |  |
| 120.0811 | 4.54 | 1.95 | 2.78 | - | ↘ |  |
| 166.0867 | 4.54 | 1.96 | 2.72 | - | ↘ |  |
| 156.0426 | 4.66 | 2.33 | 2.95 | ↘ | ↘ |  |
| 273.1216 | 4.66 | 2.38 | 2.69 | ↗ | ↘ |  |
| 104.1075 | 5.17 | 3.43 | 3.53 | ↗ | ↗ |  |
| 258.1102 | 5.17 | 3.52 | 3.57 | ↗ | ↗ | M9 [M+H]<sup>+</sup> |
| 515.2133 | 5.17 | 3.33 | 3.30 | ↗ | ↗ | M9 [2M+H]<sup>+</sup> |
| 537.1957 | 5.17 | 2.94 | 3.11 | ↗ | ↗ | M9 [2M+Na]<sup>+</sup> |
| 114.0667 | 4.24 | 2.09 | 1.89 | ↗ | - | M4 [M+H]<sup>+</sup> |
| 338.3427 | 0.73 | 2.39 | 2.24 | ↘ | ↘ | M7 [M+H]<sup>+</sup> |
| 267.2696 | 0.76 | 2.23 | 2.14 | ↗ | ↗ | M8 [M+H]<sup>+</sup>, fragment |
| 341.3062 | 0.76 | 2.39 | 2.12 | ↘ | ↘ | M8 [M+H-H<sub>2</sub>O]<sup>+</sup> |
| 359.3166 | 0.76 | 2.35 | 2.12 | ↘ | ↘ | M8 [M+H]<sup>+</sup> |
| 204.0659 | 1.28 | 2.61 | 2.67 | ↘ | ↘ | M6 [M-H]<sup>-</sup> |
| 792.6061 | 0.66 | 2.12 | 2.18 | ↘ | ↗ |  |
| 681.6038 | 0.68 | 2.02 | 1.93 | ↘ | - |  |
| 267.2696 | 0.76 | 2.23 | 2.14 | ↗ | ↗ |  |
| 341.3062 | 0.76 | 2.39 | 2.12 | ↗ | ↘ |  |
| 359.3166 | 0.76 | 2.35 | 2.12 | ↗ | ↘ |  |
| 130.1599 | 0.84 | 2.05 | 1.94 | ↘ | - |  |

|  |  |  |  |  |  |
| --- | --- | --- | --- | --- | --- |
| 168.0674 | 0.80 | 2.18 | 2.11 | ↗ | ↘ |
| 181.0982 | 1.07 | 2.49 | 2.42 | ↗ | ↗ |
| 138.0559 | 1.08 | 2.28 | 2.20 | ↗ | ↗ |
| 284.1403 | 1.15 | 2.44 | 2.42 | ↗ | ↘ |
| 167.0823 | 1.16 | 2.38 | 2.23 | ↗ | ↗ |
| 227.1397 | 1.36 | 2.43 | 2.30 | ↗ | ↗ |
| 124.0398 | 1.45 | 2.34 | 2.29 | ↗ | ↗ |
| 261.1237 | 1.46 | 2.33 | 2.12 | ↗ | ↗ |
| 197.0927 | 1.63 | 2.26 | 2.12 | ↗ | ↘ |
| 245.0957 | 1.64 | 2.00 | 2.22 | ↗ | ↘ |
| 155.0822 | 1.68 | 2.24 | 2.04 | ↗ | ↗ |
| 554.5505 | 1.76 | 0.63 | 2.00 | - | ↗ |
| 324.0593 | 3.67 | 1.78 | 2.00 | - | ↘ |
| 130.1595 | 4.81 | 2.25 | 2.14 | ↘ | ↘ |
| 160.035 | 4.80 | 2.37 | 2.19 | ↗ | ↗ |
| 296.0669 | 5.17 | 2.07 | 2.27 | ↗ | ↗ |

## References

1. Kaper JB, Nataro JP, Mobley HLT. Pathogenic Escherichia coli. Nat Rev Microbiol. févr 2004;2(2):123–40.

2. Jesudason T. WHO publishes updated list of bacterial priority pathogens. Lancet Microbe. sept 2024;5(9):100940.

3. Allocati N, Masulli M, Alexeyev MF, Di Ilio C. Escherichia coli in Europe: An Overview. Int J Environ Res Public Health. déc 2013;10(12):6235–54.

4. Breijyeh Z, Jubeh B, Karaman R. Resistance of Gram-Negative Bacteria to Current Antibacterial Agents and Approaches to Resolve It. Molecules. janv 2020;25(6):1340.

5. Husna A, Rahman MdM, Badruzzaman ATM, Sikder MH, Islam MR, Rahman MdT, et al. Extended-Spectrum β-Lactamases (ESBL): Challenges and Opportunities. Biomedicines. 30 oct 2023;11(11):2937.

6. Hu Y, Yang X, Qin J, Lu N, Cheng G, Wu N, et al. Metagenome-wide analysis of antibiotic resistance genes in a large cohort of human gut microbiota. Nat Commun. 2013;4:2151.

7. Ma Y, Liu X, Wang J. Small molecules in the big picture of gut microbiome-host cross-talk. EBioMedicine. juill 2022;81:104085.

8. Berg G, Rybakova D, Fischer D, Cernava T, Vergès MCC, Charles T, et al. Correction to: Microbiome definition re-visited: old concepts and new challenges. Microbiome. 20 août 2020;8(1):119.

9. Tang YT, He WQ. Editorial: Insights in microorganisms in vertebrate digestive systems: 2022. Front Microbiol. 5 janv 2024;14:1344969.

10. El-Sayed A, Aleya L, Kamel M. Microbiota’s role in health and diseases. Environ Sci Pollut Res Int. 2021;28(28):36967–83.

11. Huffnagle GB, Noverr MC. The emerging world of the fungal microbiome. Trends Microbiol. juill 2013;21(7):334–41.

12. Pérez JC. Fungi of the human gut microbiota: Roles and significance. Int J Med Microbiol IJMM. avr 2021;311(3):151490.

13. Maas E, Penders J, Venema K. Fungal-Bacterial Interactions in the Human Gut of Healthy Individuals. J Fungi. 19 janv 2023;9(2):139.

14. Wan X, Yang Q, Wang X, Bai Y, Liu Z. Isolation and Cultivation of Human Gut Microorganisms: A Review. Microorganisms. 20 avr 2023;11(4):1080.

15. Underhill DM, Iliev ID. The mycobiota: interactions between commensal fungi and the host immune system. Nat Rev Immunol. juin 2014;14(6):405–16.

16. Richard ML, Sokol H. The gut mycobiota: insights into analysis, environmental interactions and role in gastrointestinal diseases. Nat Rev Gastroenterol Hepatol. 2019;16(6):331–45.

17. Mukherjee PK, Sendid B, Hoarau G, Colombel JF, Poulain D, Ghannoum MA. Mycobiota in gastrointestinal diseases. Nat Rev Gastroenterol Hepatol. févr 2015;12(2):77–87.

18. Hoffmann C, Dollive S, Grunberg S, Chen J, Li H, Wu GD, et al. Archaea and fungi of the human gut microbiome: correlations with diet and bacterial residents. PloS One. 2013;8(6):e66019.

19. Catalán-Serra I, Thorsvik S, Beisvag V, Bruland T, Underhill D, Sandvik AK, et al. Fungal Microbiota Composition in Inflammatory Bowel Disease Patients: Characterization in Different Phenotypes and Correlation With Clinical Activity and Disease Course. Inflamm Bowel Dis. 16 déc 2023;30(7):1164–77.

20. Nobile CJ, Johnson AD. Candida albicans Biofilms and Human Disease. Annu Rev Microbiol. 2015;69:71–92.

21. Hager CL, Ghannoum MA. The mycobiome: Role in health and disease, and as a potential probiotic target in gastrointestinal disease. Dig Liver Dis. 1 nov 2017;49(11):1171–6.

22. Peters BM, Jabra-Rizk MA, O’May GA, Costerton JW, Shirtliff ME. Polymicrobial Interactions: Impact on Pathogenesis and Human Disease. Clin Microbiol Rev. janv 2012;25(1):193–213.

23. Deveau A, Bonito G, Uehling J, Paoletti M, Becker M, Bindschedler S, et al. Bacterial– fungal interactions: ecology, mechanisms and challenges. FEMS Microbiol Rev. 1 mai 2018;42(3):335–52.

24. Eshima S, Kurakado S, Matsumoto Y, Kudo T, Sugita T. Candida albicans Promotes the Antimicrobial Tolerance of Escherichia coli in a Cross-Kingdom Dual-Species Biofilm. Microorganisms. nov 2022;10(11):2179.

25. Sokol H, Leducq V, Aschard H, Pham HP, Jegou S, Landman C, et al. Fungal microbiota dysbiosis in IBD. Gut. juin 2017;66(6):1039–48.

26. Want EJ, Wilson ID, Gika H, Theodoridis G, Plumb RS, Shockcor J, et al. Global metabolic profiling procedures for urine using UPLC-MS. Nat Protoc. juin 2010;5(6):1005–18.

27. Boccard J, Rudaz S. Harnessing the complexity of metabolomic data with chemometrics. J Chemom. 2014;28(1):1–9.

28. Qannari EM, Wakeling I, Courcoux P, MacFie HJH. Defining the underlying sensory dimensions. Food Qual Prefer. 1 janv 2000;11(1):151–4.

29. Rosa LN, de Figueiredo LC, Bonafé EG, Coqueiro A, Visentainer JV, Março PH, et al. Multi-block data analysis using ComDim for the evaluation of complex samples: Characterization of edible oils. Anal Chim Acta. 8 avr 2017;961:42–8.

30. Thévenot EA, Roux A, Xu Y, Ezan E, Junot C. Analysis of the Human Adult Urinary Metabolome Variations with Age, Body Mass Index, and Gender by Implementing a Comprehensive Workflow for Univariate and OPLS Statistical Analyses. J Proteome Res. 7 août 2015;14(8):3322–35.

31. Wiklund S, Johansson E, Sjöström L, Mellerowicz EJ, Edlund U, Shockcor JP, et al. Visualization of GC/TOF-MS-Based Metabolomics Data for Identification of Biochemically Interesting Compounds Using OPLS Class Models. Anal Chem. 1 janv 2008;80(1):115–22.

32. Schymanski EL, Jeon J, Gulde R, Fenner K, Ruff M, Singer HP, et al. Identifying Small Molecules via High Resolution Mass Spectrometry: Communicating Confidence. Environ Sci Technol. 18 févr 2014;48(4):2097–8.

33. Selegato DM, Castro-Gamboa I. Enhancing chemical and biological diversity by co-cultivation. Front Microbiol. 2023;14:1117559.

34. De Brucker K, Tan Y, Vints K, De Cremer K, Braem A, Verstraeten N, et al. Fungal β-1,3-Glucan Increases Ofloxacin Tolerance of Escherichia coli in a Polymicrobial E. coli/Candida albicans Biofilm. Antimicrob Agents Chemother. 14 mai 2015;59(6):3052–8.

35. Farrokhi Y, Al-shibli B, Al-hameedawi DFJ, Neshati Z, Makhdoumi A. Escherichia coli enhances the virulence factors of Candida albicans, the cause of vulvovaginal candidiasis, in a dual bacterial/fungal biofilm. Res Microbiol. 1 juin 2021;172(4):103849.

36. Yang W, Zhou Y, Wu C, Tang J. Enterohemorrhagic Escherichia coli promotes the invasion and tissue damage of enterocytes infected with Candida albicans in vitro. Sci Rep. 22 nov 2016;6:37485.

37. Thein ZM, Samaranayake YH, Samaranayake LP. Effect of oral bacteria on growth and survival of Candida albicans biofilms. Arch Oral Biol. 1 août 2006;51(8):672–80.

38. Park SJ, Han KH, Park JY, Choi SJ, Lee KH. Influence of bacterial presence on biofilm formation of Candida albicans. Yonsei Med J. mars 2014;55(2):449–58.

39. Cabral DJ, Penumutchu S, Norris C, Morones-Ramirez JR, Belenky P. Microbial competition between Escherichia coli and Candida albicans reveals a soluble fungicidal factor. Microb Cell Graz Austria. 7 mars 2018;5(5):249–55.

40. Bandara HMHN, Yau JYY, Watt RM, Jin LJ, Samaranayake LP. Escherichia coli and its lipopolysaccharide modulate in vitro Candida biofilm formation. J Med Microbiol. déc 2009;58(Pt 12):1623–31.

41. Mason KL, Erb Downward JR, Mason KD, Falkowski NR, Eaton KA, Kao JY, et al. Candida albicans and Bacterial Microbiota Interactions in the Cecum during Recolonization following Broad-Spectrum Antibiotic Therapy. Infect Immun. 11 sept 2012;80(10):3371–80.

42. Stahl PD, Klug MJ. Characterization and differentiation of filamentous fungi based on Fatty Acid composition. Appl Environ Microbiol. nov 1996;62(11):4136–46.

43. Stenz L, François P, Fischer A, Huyghe A, Tangomo M, Hernandez D, et al. Impact of oleic acid (cis-9-octadecenoic acid) on bacterial viability and biofilm production in Staphylococcus aureus. FEMS Microbiol Lett. oct 2008;287(2):149–55.

44. Vascellari S, Palmas V, Melis M, Pisanu S, Cusano R, Uva P, et al. Gut Microbiota and Metabolome Alterations Associated with Parkinson’s Disease. mSystems. 15 sept 2020;5(5):e00561–20.

45. Arora D, Gupta P, Jaglan S, Roullier C, Grovel O, Bertrand S. Expanding the chemical diversity through microorganisms co-culture: Current status and outlook. Biotechnol Adv. 2020;40:107521.

